# A novel *Enterococcus faecalis* heme transport regulator (FhtR) is a heme sensor in the gastrointestinal tract

**DOI:** 10.1101/2020.06.22.166298

**Authors:** Vincent Saillant, Damien Lipuma, Emeline Ostyn, Laetitia Joubert, Alain Boussac, Hugo Guerin, G. Brandelet, P. Arnoux, Delphine Lechardeur

## Abstract

*Enterococcus faecalis* is a commensal Gram-positive pathogen found in the intestines of mammals, and is also a leading cause of severe infections occurring mainly among antibiotic-treated dysbiotic hospitalized patients. Like most intestinal bacteria, *E. faecalis* does not synthesize heme. Nevertheless, environmental heme can improve *E. faecalis* fitness by activating respiration metabolism and a catalase that limits hydrogen peroxide stress. Since free heme also generates redox toxicity, its intracellular levels need to be strictly controlled. Here, we describe a unique transcriptional regulator, FhtR, (<u>F</u>aecalis <u>h</u>eme <u>t</u>ransport <u>R</u>egulator), which manages heme homeostasis by controlling an HrtBA-like efflux pump (named HrtBA_*Ef*_). We show that FhtR, by managing intracellular heme concentration, regulates the functional expression of the heme dependent catalase A (KatA), thus participating in heme detoxification. The biochemical features of FhtR binding to DNA, and its interactions with heme that induce efflux, are characterized. The FhtR-HrtBA_*Ef*_ system is shown to be relevant in a mouse intestinal model. We further show that FhtR senses heme from blood and hemoglobin but also from crossfeeding by *Escherichia coli*. These findings bring to light the central role of FhtR heme sensing in response to heme fluctuations within the gastrointestinal tract, which allow this pathogen to limit heme toxicity while ensuring expression of an oxidative defense system.

**Importance:** *Enterococcus faecalis*, a normal, harmless colonizer of the human intestinal flora can cause severe infectious diseases in immunocompromised patients, particularly those that have been heavily treated with antibiotics. Therefore, it is important to understand the factors that promote its resistance and its virulence. Here, we report a new mechanism used by *E. faecalis* to detect the concentration of heme, an essential but toxic metabolite that is present in the intestine. *E. faecalis* needs to scavenge this molecule to respire and fight stress generated by oxydants. Heme sensing triggers the synthesis of a heme efflux pump that balances the amount of heme inside the bacteria. With this mechanism, *E. faecalis* can use heme without suffering from its toxicity.

## Introduction

*Enterococcus faecalis* is a commensal inhabitant of the gastrointestinal tract (GIT) and a subdominant species in the core intestinal microbiota of healthy humans and other mammals (1). This lactic acid bacterium is also a major opportunistic pathogen that causes a large number of nosocomial infections such as endocarditis, bacteremia, urinary tract infections or meningitis (2). In recent decades, *E. faecalis* has emerged as a leading cause of enterococcal infections (70 %), and is the third most frequent source of hospital-acquired nosocomial infections. *E. faecalis* is thus considered as a major public health threat due to its intrinsic resistance to antibiotics and emergence of multidrug-resistant isolates (3). Selective outgrowth of enterococci following intestinal dysbiosis is frequent, regardless of whether dysbiosis results from antibiotic treatment, intestinal inflammation, or infection (4). In addition to intrinsic and acquired antibiotic resistances, *E. faecalis* is resistant to other antimicrobial factors, such as bile, and tolerates a wide variety of stress factors such as temperature, pH, oxygen tension, or oxidation (1).

For most living organisms, heme^1^ (iron porphyrin) is an essential cofactor of enzymes such as cytochromes, catalases or peroxidases (5). The importance of heme resides in the unique properties of its iron center, including the capacity to undergo electron transfer, perform acidbase reactions, and interact with various coordinating ligands (6). Paradoxically, the high potential redox of heme explains its toxicity (7, 8). Most bacteria carry the enzymatic machinery for endogenous heme synthesis. However, numerous bacteria, designated as heme-auxotrophs, lack some or all the enzymes needed for autosynthesis, but still require this molecule for their metabolism (5). Interestingly, *E. faecalis* is a heme-auxotroph, like the majority of species constituting the gastrointestinal microbiota (9, 10). When heme is added to an aerated culture, *E. faecalis* activates a terminal cytochrome *bd* oxidase, causing a shift from fermentation to an energetically favorable respiratory metabolism (10, 11). *E. faecalis*, unlike other heme-auxotrophic Firmicutes, also encodes a heme catalase, (KatA; EC 1.11.1.6), limiting hydrogen peroxide stress when heme is available (9). Both functions contribute to the virulence of several Gram positive pathogens (12, 13). Although the importance of heme as a cofactor for numerous cellular functions is established (5, 14), the mechanisms governing exogenous heme internalization and secretion wich contribute to heme homeostasis vary among bacteria and are poorly understood. In contrast, heme homeostasis must be strictly regulated in all bacteria to avoid toxicity (6, 8). Heme efflux is a documented defence mechanism against heme toxicity in some Firmicutes: i) the Pef regulon comprises 2 multi-drug resistance efflux pumps and a MarR-type heme-responsive regulator in *Streptococcus agalactiae* (15); ii) the HatRT system comprises a TetR family heme binding transcriptional regulator (*hatR*) and a major facilitator superfamily heme transporter (*hatT*) in *Clostridium difficile* (16); iii) Heme homeostasis in several Gram positive bacteria relies on Hrt (heme-regulated transport) proteins HrtB (permease) and HrtA (ATPase), an ABC transporter, which promote heme efflux (13, 17–19). Expression of *hrtBA* is controlled by *hssRS* genes, encoding a two-component heme sensor and response regulator in numerous Gram positive pathogens including *S. agalactiae, Staphylococcus aureus* and *Bacillus anthracis*, (13, 18–21). In contrast, the food bacterium *Lactococcus lactis* regulates HrtBA expression *via* the TetR family heme sensor (HrtR) (18). To date, the mechanisms involved in *E. faecalis* management of environmental heme are unknown.

In this work, we describe the mechanism by which a novel *E. faecalis* TetR regulator, called FhtR (<u>F</u>aecalis <u>h</u>eme <u>t</u>ransporter <u>r</u>egulator), induces expression of HrtBA_*Ef*_, a conserved heme effux transporter. We show that FhtR binds intracellular heme resulting in de-repression and increased transcription of *hrtBA_Ef_*. Heme iron coordination specifies FhtR as a heme sensor and we defined a critical role for the tyrosine 132. Our results also establish this system as a master mechanism of control of intracellular heme availability as shown by its requirement for the expression of the heme dependent *E. faecalis* KatA. Our conclusions lead to a new picture of heme homeostasis in *E. faecalis*. Finally, the relevance of the FhtR system to *E. faecalis* is shown in a mouse intestine model, suggesting the importance of FhtR for gastrointestinal lumen heme homeostasis.

## Results

### The conserved heme efflux transporter, HrtBA_*Ef*_, is functional in *E. faecalis*

*E. faecalis* OG1RF genome encodes two adjacent ORFs, OG1RF_RS02770 and OG1RF_RS02775 sharing respectively 24 % and 45 % amino-acid (AA) sequence identity with HrtB and HrtA from *Staphylococcus aureus* (17) (Fig. S1). We thus verified the role of these ORFs, referred to as HrtB_*Ef*_ and HrtA_*Ef*_, in *E. faecalis* heme efflux. Compared to the wild type (WT) OG1RF strain, growth of an in-frame Δ*hrtBA_Ef_* deletion mutant was severely impaired at hemin concentrations ≥ 10 μM, highlighting the involvement of HrtBA_*Ef*_ in limiting heme toxicity in *E. faecalis* (Fig. 1A). The Δ*hrtBA_Ef_* mutant grown in 5 μM heme-containing medium accumulated about twice more intracellular heme than the WT strain, as evaluated by the pyridine hemochrome assay (22) (Fig. 1B). This result correlated with the intense red color of culture pellets from the Δ*hrtBA_Ef_* mutant compared to the WT strain (Fig. 1C). Intracellular heme concentrations were also followed using the intracytoplasmic heme sensor HrtR (18): β–gal activity from the reporter plasmid P_hrt_-*hrtR*-*lac*, was about 4 times higher in Δ*hrtBA_Ef_* compared to the WT exposed to 5 μM heme (Fig. 1D). Thus, *E. faecalis* expresses a functional HrtBA_*Ef*_ heme efflux transporter that modulates intracellular heme levels.

**Figure 1.**
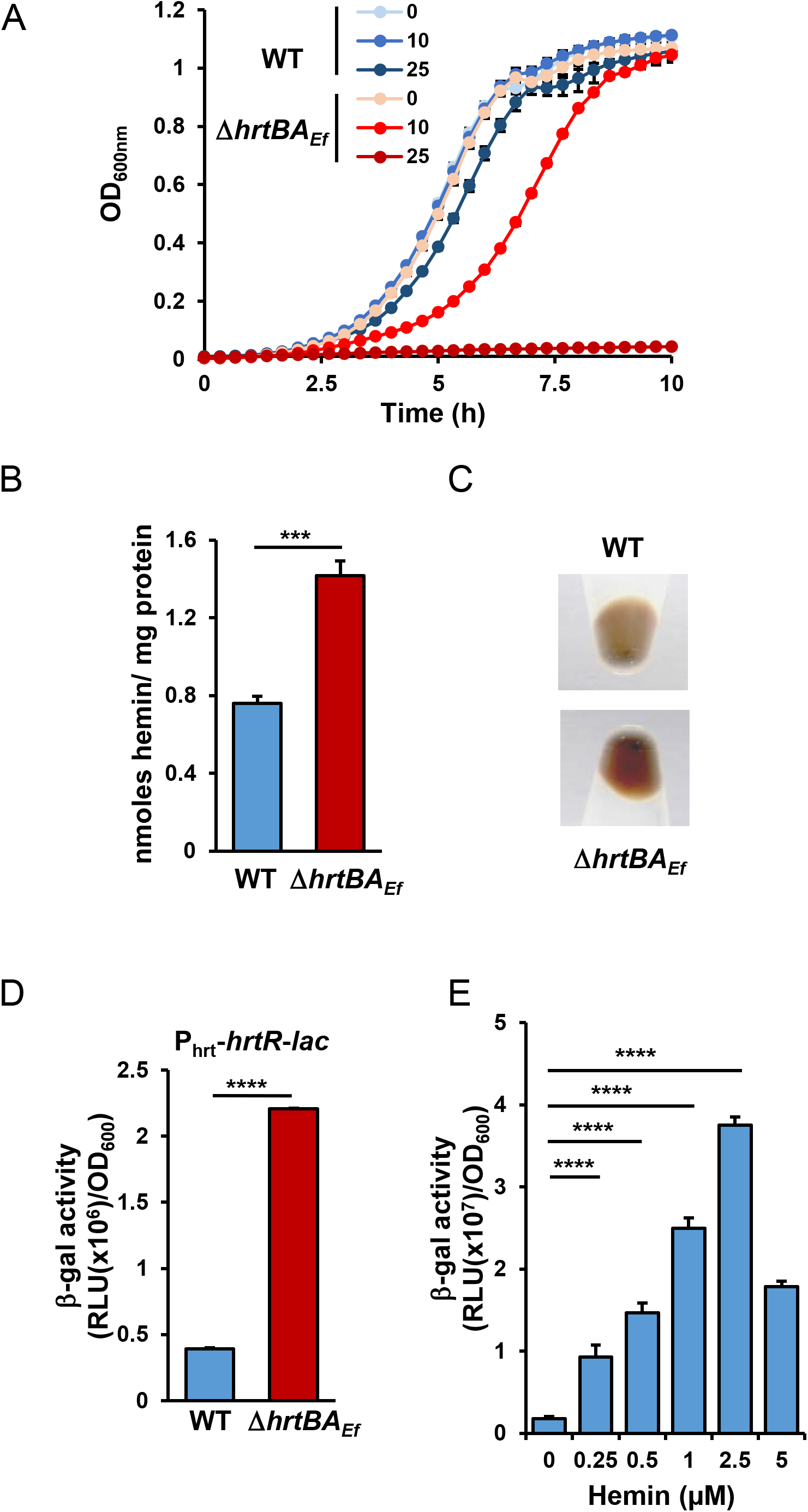
HrtBA_*Ef*_ controls and responds to heme intracellular cellular content in *E. faecalis*. **(A)** Deletion of *hrtBA_Ef_* increases sensitivity to hemin toxicity. Overnight cultures of WT and Δ*hrtBA_Ef_* strains were diluted to OD_600nm_=0.01 and grown with the indicated concentrations of hemin (μM) for 10 h at 37 °C in a microplate Spark spectrophotometer (Tecan). OD_600nm_ was measured every 20 min. Results represent the average ± S.D. from triplicate technical samples. Result is representative of 3 independent experiments. **(B)** Heme accumulates in Δ*hrtBA_Ef_* strain. WT and Δ*hrtBA_Ef_* strains were grown to OD_600nm_=0.5 prior addition of 5 μM hemin in the culture medium for an additional 1.5 h. Bacteria were pelleted by centrifugation and heme content was determined by the pyridine hemochrome assay on cell lysates. Heme content was normalized to the protein concentration. Background from bacteria non-incubated with hemin was substracted. Results represent the average ± S.D. from biological triplicates. ***, *P* = 0.0002, Student’s *t* test. **(C)** HrtBA_*Ef*_ participates in the color phenotype of *E. faecalis* upon heme exposure. Cells were grown as in (A), were incubated for 1.5 h with 5 μM hemin. The bacteria were photographed following centrifugation. Result is representative of 3 independent experiments. **(D)** HrtBA_*Ef*_ reduces heme cytoplasmic concentration. WT and Δ*hrtBA_Ef_* strains carrying the intracellular sensor plasmid, pP_hrt_-*hrtR*-*lac* were grown as in (B). B-gal activity was quantified by luminescence after 1.5 h incubation with 5 μM hemin. Results represent the average ± S.D. from triplicate technical samples. ****, *P* < 0.0001, Student’s *t* test. Values with no added hemin are substracted. Result is representative of 3 independent experiments. **(E)** Induction of *hrtBA_Ef_* operon by hemin. WT OG1RF strain transformed with the reporter plasmid pP_hrtBA_-*lac* was grown and B-gal activity was determined as in (D) following incubation with the indicated concentrations of hemin. Results represent the average ± S.D. from triplicate technical samples. ****, *P* <0.001, one-way ANOVA followed by Dunnett’s post test comparison to 0 μM. Result is representative of 3 independent experiments.

Transcriptional regulation of *hrtBA_Ef_* by heme was then investigated using a *hrtBA_Ef_* promoter reporter, P_hrtBA_-*lac*. β–gal expression in the WT strain was induced by 0.1 to 2.5 μM hemin in the culture medium. Induction reached a maximum at concentrations below 5 μM (Fig. 1E). This concentration range is far below WT strain sensitivity to heme toxicity (≥ 25 μM) (Fig. 1A). We conclude that HrtBA_*Ef*_ expression is induced at subtoxic heme concentrations.

### A new TetR-regulator, FhtR, controls *hrtBA_Ef_* expression

The above findings prompted us to investigate the mechanism of *hrtBA_Ef_* induction. Several Gram positive pathogens regulate *hrtBA_Ef_ via* an adjacent 2 component system HssR and HssS (13, 19, 20). No *hssR hssS* genes were identified in near the *hrtBA_Ef_* operon in OG1RF or other *E. faecalis* genomes. However, a monocistronic gene encoding a TetR family transcriptional regulator, OG1RF_RS02765, is adjacent to *hrtBA_Ef_*(Fig 2A), sharing no significant AA identity with the *hrtBA* regulator, HrtR in *Lactococcus lactis* (18). We hypothesized that OG1RF_RS02765 was the transcriptional regulator of *hrtBA_Ef_* and tentatively named it FhtR (<u>F</u>aecalis <u>h</u>eme <u>t</u>ransport <u>R</u>egulator) (Fig. 2A).

**Figure 2.**
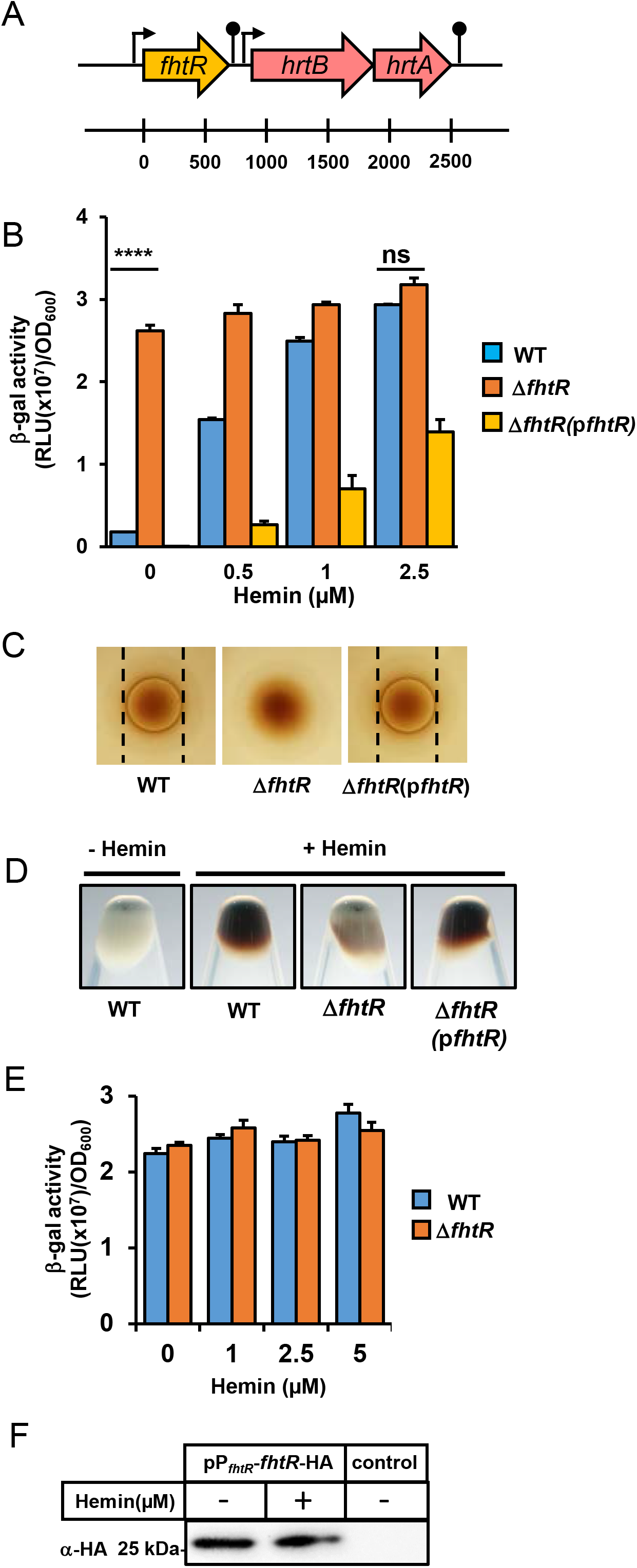
FhtR, a novel heme sensing transcriptional regulator controls *hrtBA_Ef_* expression in *E. faecalis*. **(A)** Schematic representation of the *fhtR* gene and concomitant *hrtBA_Ef_* operon. The *fhtR* gene (OG1RF_RS02765) encodes a TetR family transcriptional regulator. The *hrtB_Ef_* (OG1RF_RS02770) and *hrtA_Ef_* (OG1RF_RS02775) genes encode a permease and ATPase, respectively. **(B)** FhtR controls *hrtBA_Ef_* expression. WT and Δ*fhtR* strains carrying the reporter plasmid pP_hrtBA_-*lac*, and Δ*fhtR* carrying a plasmid, p*fhtrR*, combining both P_hrtBA_-*lac* and P_fhtR_-*fhtR* were grown to OD_600nm_=0.5 and Bgal expression was quantified by luminescence as reported in Fig. 1 with the indicated concentrations of hemin. Results represent the average ± S.D. from triplicate technical samples. ****, *P* < 0.001; ns, non significant, Student’s *t* test. Result is representative of 3 independent experiments **(C)** Visualization of the impact of FhtR on heme accumulation. WT, Δ*fhtR* and Δ*fhtR* complemented with the expression plasmid p*fhtR* were grown and incubated with 5 μM hemin as in (A). The cell were pelleted by centrifugation and photographed. Result is representative of 3 independent experiments. **(D)** *fhtR* deletion abrogates heme toxicity. Stationary phase cultures of WT, Δ*fhtR* and Δ*fhtR*(p*fhtR*) were plated on solid medium. Hemin (10 μl of a 10 mM stock solution) was pipetted directly onto plates, which were incubated for 24 h. Inhibition zones appear as dark clearing in the center of each panel. No inhibition zone was observed for Δ*fhtR* strain. Result is representative of 3 independent experiments. **(E)** FhtR expression is constitutively induced. Bgal expression upon hemin addition to the culture medium in WT and Δ*fhtR* strains transformed with the pPfhtr-lac reporter plasmid was determined by luminescence as in Fig. 1. **(F)** *fhtR* transcription is not mediated by hemin. Δ*fhtR* strain transformed with pP_fhtR_-*fhtR*-HA or empty vector (control) was used to follow FhtR expression by Western Blot using an anti-HA antibody. Bacteria were grown to OD_600nm_= 0.5 and incubated with 2.5 μM hemin for 1.5 h. SDS-PAGE was performed on cell lysates (80 μg per lane).

To investigate the role of FhtR in heme-dependent transcription of *hrtBA_Ef_*, an *fhtR* inframe deletion in OG1RF (Δ*fhtR*) was constructed and transformed either with pP_hrtBA_-*lac* or p*fhtR* encompassing both (pP_hrtBA_-*lac* and P_fhtR_-*fhtR*) expression cassettes (Fig. 2B). In contrast to the WT strain, β–gal was expressed independently of heme in Δ*fhtR*(pP_hrtBA_-*lac*) (Fig. 2B). Complementation by p*fhtR* partially restored hemin-dependent expression compared to the WT strain (Fig. 2B). Moreover, on solid medium, Δ*fhtR* exhibited a complete absence of sensitivity to heme compared to WT and complemented Δ*fhtR*(p*fhtR*) strains (Fig. 2C). These results are in line with the observation that heme accumulation is reduced in Δ*fhtR* compared to the WT or Δ*fhtR*(p*fhtR*) strains (Fig. 2D), and that HrtBA_*Ef*_ may be constitutively expressed in the Δ*fhtR* mutant (Fig. 2B). P_fhtR_ was constitutively transcribed, with no effects of heme, nor of FhtR expression as shown using P_fhtR_-*lac* as reporter (Fig. 2E), and by Western Blot (WB) using FhtR-HA tagged fusion expressed from P_fhtR_ (Fig. 2F). We conclude that *E. faecalis* uses a constitutively expressed, unique intracellular heme sensor, FhtR, to control *hrtBA_Ef_* expression.

### FhtR is a heme binding protein

Members of the TetR family of transcriptional regulators act as chemical sensors (23, 24). Ligand binding alleviates TetR protein interactions with their respective operators, leading to promoter induction (23, 24). To verify that heme was the signal that relieves FhtR-mediated *hrtBA_Ef_* repression, recombinant FhtR was purified as a fusion to the maltose binding protein (MBP-FhtR) from *Escherichia coli*. MBP-FhtR appeared green (Fig. 3A, inset) and its UV-visible spectrum exhibited a strong Soret band suggesting that FhtR scavenges the endogenously produced heme (Fig. 3A). To purify an apoFhtR, the MBP-FhtR was expressed from a heme synthesis deficient *E. coli* strain (*hemA::kan*) (Fig. 3B, dashed line). The purified protein bound hemin *b in vitro* (*i.e*., non-covalently) with a similar UV-visible spectrum as observed above for *in vivo* bound heme: a Soret band at 407 nm and Q bands at 491 nm, 528 nm and 608 nm (Fig. 3B, black line). Size-exclusion chromatography profiles showed that both apo- and holo-MBP-FhtR eluted as a single pea corresponding to the size of a dimer (132 kDa), in line with other TetR regulators (Fig. 3C) (23). The 608 nm charge transfer band and Soret at 407 nm are indicative of a ferric high-spin tyrosinate-ligated heme where heme is anchored through a proximal tyrosinate side chain (25, 26). Hemin pentacoordinate high-spin ligation to FhtR was further confirmed by EPR spectroscopy (see below). Heme dissociation coefficient (*K*_d_) was 310 nM as determined by MBP-FhtR fluorescence quenching over increasing concentration of hemin (Fig. 3D). Heme titration by differential absorption spectroscopy at 407 nm showed that the saturation point corresponded to the binding of one molecule of hemin per MBP-FhtR monomer (Fig. 3D, inset). Altogether, these data demonstrate that FhtR is a heme-binding protein, suggesting that heme interaction is the primary event leading to activation of *hrtBA_Ef_* transcription.

**Figure 3.**
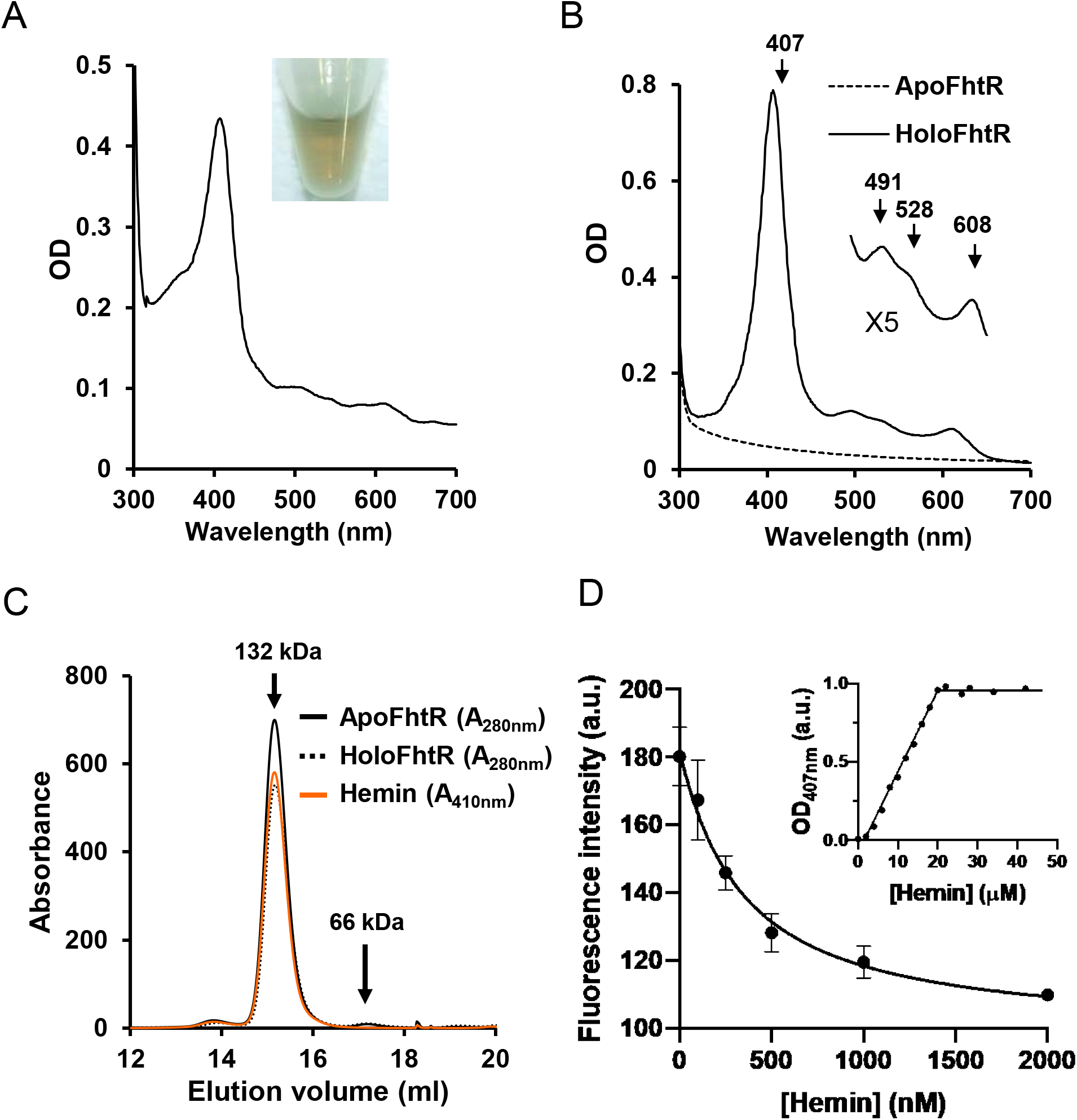
FhtR binds heme. **(A)** UV-visible absorption spectra of MBP-FhtR as purified from *E. coli*. UV-visible spectra of 30 μM (in 200 μl) MBP-FhtR was obtained in a microplate spectrophotometer (Spark, Tecan) and normalized to OD_280nm_=1. *Inset*, a photograph of the purified MBP-FhtR. Results are representative of 3 experiments. **(B)** UV-visible spectra of apoMBP-FhtR complexed with hemin. MBP-FhtR was purified from *E. coli* (*hemA::kan*) strain as an apoprotein (dashed line) that was mixed with equimolar concentration of hemin. Spectra was obtained as in (A) with 20 μM complex and normalized to OD_280nm_=1. Inset, magnification of the 500-700 nm region. Results are representative of 3 experiments. **(C)** Size-exclusion chromatography of apo- and holoMBP-FhtR. MBP-FhtR was purified and complexed with hemin as in (B). 40 μM in 100 μl were loaded on a Superdex 200 Increase 10/300 GL gel filtration column (GE healthcare) in HEPES 20 mM NaCl 300 mM buffer. Protein and heme content were analyzed at OD_280nm_ and OD_410nm_ respectively. Results are representative of 3 experiments. **(D)** Titration of MBP-FhtR with hemin followed by fluorescence and absorbance (inset). For the fluorescence experiment, 50 nM of ApoMBP-FhtR purified from (*hemA*::*kan*) *E. coli* as in (B) were titrated with increasing increments of hemin. Fluorescence intensity was recorded and plotted against hemin concentration. The experiment was repeated three times, fitted using the nonlinear regression function of GraphPad Prism 4 software and gave a *K*_d_ of 310 nM. The inset depicts the absorbance at 407_nm_ of ApoMBP-FhtR plotted against hemin concentration. Curve is representative of 10 independent experiments and was fitted using the nonlinear regression function of GraphPad Prism 4 software and determined that the stoichiometry of the FhtR:hemin complex was 1:1.

### Tyrosine 132 is a crucial heme axial ligand in FhtR

According to UV-visible and EPR spectra, the likely candidate for axial ligand of oxidized heme is a tyrosine (Y) (see above), several Y residues present in FhtR were substituted to phenylalanine (F) (Fig. S2A). F and Y both have phenyl ring structures, so that F substitution minimizes an impact on FhtR conformation. Although, F lacks the hydroxyl group that coordinates heme. FhtR heme binding was not modified for several mutants tested individually (in blue, Fig. S2A) except, FhtR^Y132F^, which was purified from *E. coli* with a strong decrease in heme content compared to WT MBP-FhtR, indicating a loss of heme affinity *in vivo* (Fig. 4A). Surprisingly, apoMBP-FhtR^Y132F^ purified from *hemA::kanA E. coli*; Fig. 4B) exhibited similar UV-visible spectra and *K_d_* upon hemin addition (data not shown), questioning the implication of this tyrosine in heme binding. The role of Y132 was further analysed by EPR (Electron Paramagnetic Resonance) spectroscopy (Fig. 4C). HoloMBP-FhtR exhibited an axial high spin (*S* = 5/2) heme signal with two well-defined resonances at around *g* ~ 6 (with a crossing point at 1190 gauss) (Fig. 4C, inset) and a resonance at *g* ~ 2 (~ 3390 gauss), indicative of a 5-coordinated Fe^III^ structure. Although the UV-visible spectra of FhtR and FhtR^Y132F^ supplemented with hemin do not differ to a detectable level, the EPR spectra of FhtR^Y132F^ was significantly modified thus showing that either the ligand of the iron has been exchanged for another one or more likely, the interaction of the axial ligand with its environment has been modified. To conciliate these results, it is possible that while Y132 is the primary ligand, another distal ligand can take over ligation in the Y132F mutant to become the dominant ligand *in vitro* (meanwhile hydrophobic contacts would insure retention of the binding affinity).

**Figure 4.**
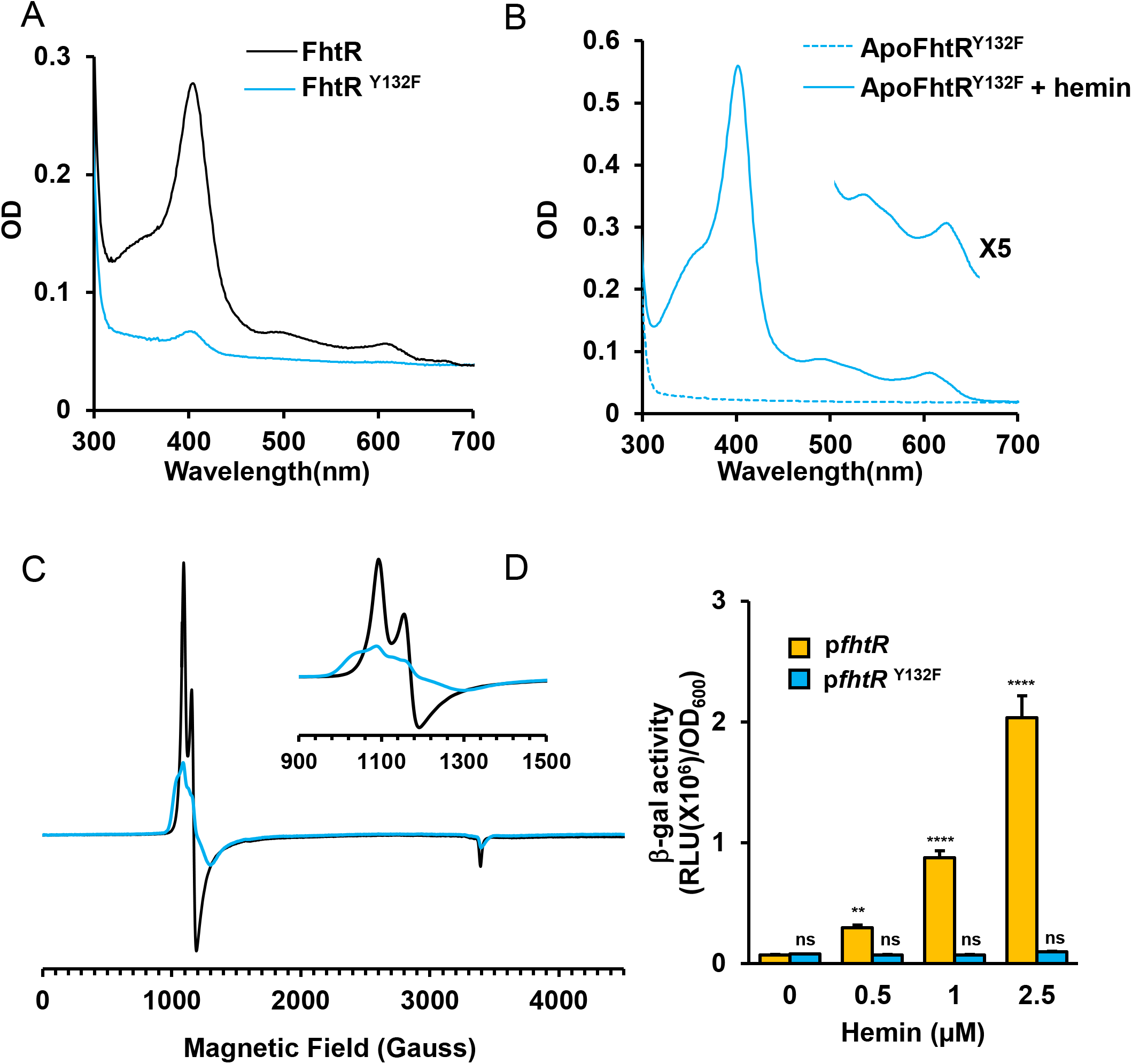
Ligation of FhtR with hemin implicates the tyrosine Y132. **(A)** Comparative UV-visible absorption spectra of 20 μM MBP-FhtR and MBP-FhtR^Y132F^ purified from *E. coli*. UV-visible spectra was performed as in Fig. 3A. **(B)** UV-visible spectra of apoMBP-FhtR^Y132F^ complexed with hemin. MBP-FhtR^Y132F^ was purified from *E. coli* (*hemA*::kan) strain as an apoprotein (dashed line) that was mixed with equimolar concentration of hemin (holoMBP-FhtR). Spectra was obtained as in (A) with 20 μM complex and normalized to OD_280nm_ = 1. Inset, magnification of the 500-700 nm region. Results are representative of 3 experiments. **(C)** EPR spectroscopy of MBP-FhtR and MBP-FhtR^Y132F^ in complex with hemin. EPR spectra of 60 μM bound hemin to WT (black line) and Y132 mutant (red line) MBP-FhtR (red line) in 20 mM HEPES, pH 7; 300 mM NaCl were recorded. Inset represents a magnification of the 900-1500 Magnetic Field range. **(D)** Induction of *hrtBA_Ef_* operon by hemin. Bgal activity from Δ*fhtR* OG1RF strain transformed either with pP_hrtBA_-*lac*; P_fhtR_-*fhtR* or pP_hrtBA_-*lac*; P_fhtR_-fhtR^Y132F^ was determined as in Fig. 1C following incubation with the indicated concentrations of hemin. Results represent the average ± S.D. from triplicate technical samples. **, P= 0.0058, ****, *P* < 0.0001, ns, non significant, one-way ANOVA followed by Dunnett’s posttest comparison to WT (0 μM). Result is representative of 3 independent experiments.

We then compared FhtR and FhtR^Y132F^ activities *in vivo*. The Δ*fhtR* mutant was complemented either with p*fhtR* (pP_hrtBA_-*lac*; P_fhtR_-*fhtR*) or p*fhtR*^Y132F^ (pP_hrtBA_-*lac*; P_fhtR_-*fhtR*^Y132F^) and β–gal expression was followed upon hemin addition to medium (Fig. 4D). Expression of both proteins prevented *hrtBA_Ef_* transcription in the absence of heme, in contrast to full expression in Δ*fhtR* (Fig. 4D). However, hemin addition led to P_hrtBA_-*lac* expression in the strain carrying WT FhtR, but not FhtR^Y132F^. Taking into account that WT FhtR and FhtR^Y132F^ were expressed to similar levels as confirmed on WB (Fig. S2B), impaired derepression of P_hrtBA_ by FhtR^Y132F^ in the presence of heme further implicates Y132 at least, one of the main proximal ligand. Altogether, these data specify FhtR Y132 as a critical residue in the coordination of heme with FhtR, which enables *hrtBA_Ef_* transcription.

### FhtR controls *hrtBA_Ef_* transcription by binding 2 distinct 14-nt palindromic repeat sequences

TetR family operators usually comprise a 10-30 nucleotides (nt) inverted repeat sequence with internal palindromic symmetry (23). Two such 14-nt long palindromes were identified in the −10/-35 region of the *hrtBA_Ef_* promoter (called P1 and P2; Fig. 5A). An electrophoretic mobility shift assay (EMSA) was performed with apoMBP-FhtR, using a 325-bp DNA segment comprising the *hrtBA_Ef_* promoter (Fig. 5B), or a segment covering the internal *hrtB* region as control (Fig. S3). FhtR specific interaction with the P_*hrtBA*_ DNA segment confirmed FhtR binding specificity. The shifted DNA migrated as 2 distinct bands (C1 and C2), in proportions that depended on the MBP-FhtR: DNA ratio (Fig. 5B), plausibly revealing that FhtR complexes with either 1 or 2 palindromes (Fig. 5B). To test this, we performed random substitutions of P1 and/or P2 nucleotides (P1* and P2*) and analyzed DNA shifts with EMSA. Replacement of both distal and proximal operators (P_hrtBA P1*, P2*_) abolished the FhtR induced DNA shift, confirming the role of palindromes in the interaction of FhtR with P_hrtBA_ (Fig. 5C). Single replacement of P1 (P_hrtBA P1*_) or P2 (P_hrtBA P2*_) resulted in complete DNA shifts that migrated faster in the gel (C1) than seen with the native nucleotide sequence (Fig. 5C). We conclude that both P1 and P2 are FhtR binding sites.

**Figure 5.**
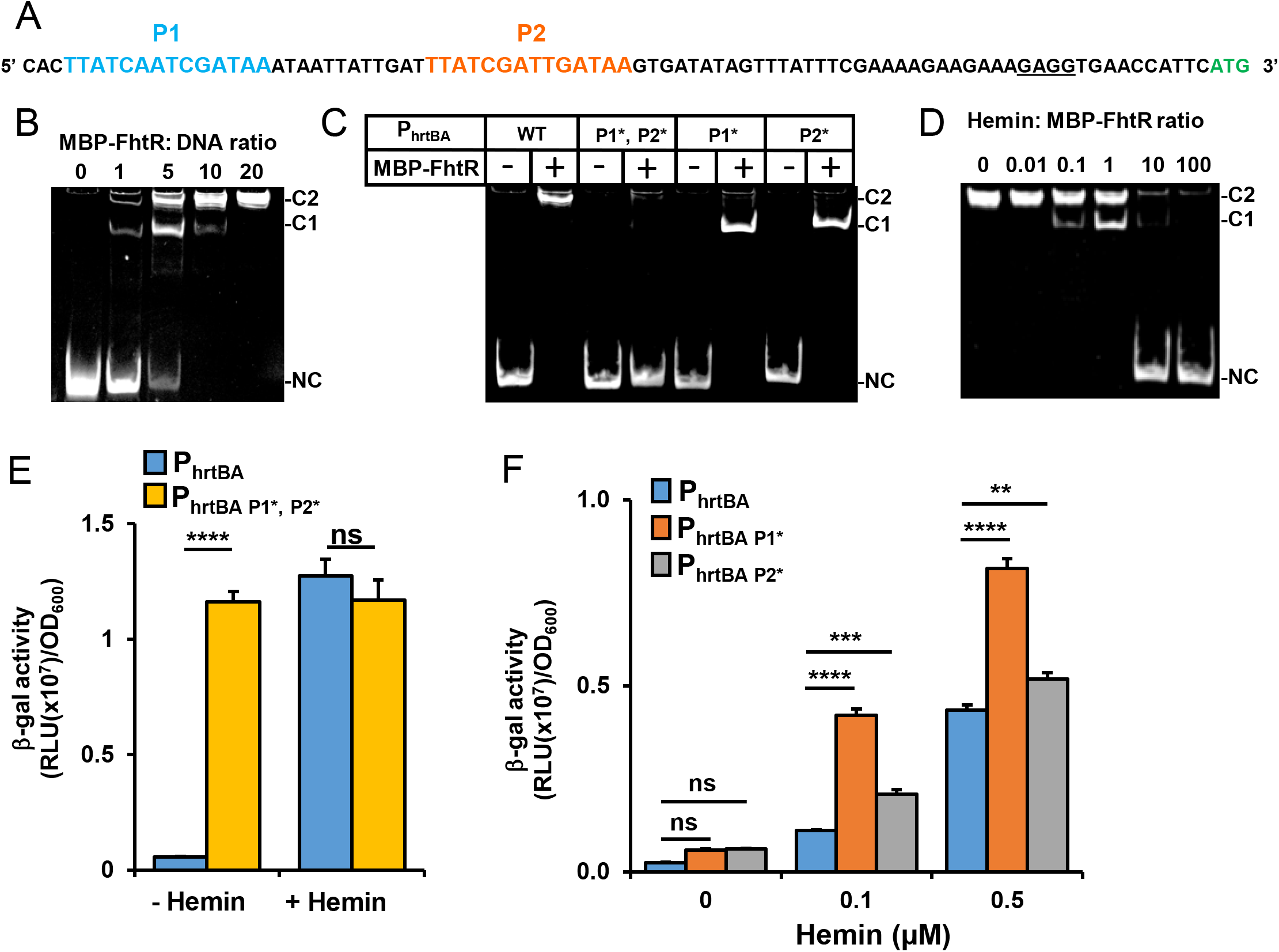
FhtR controls *hrtB_Ef_* transcription via binding to 2 repeated 14-nt palindromic sequences. **(A)** Two 14 nucleotides palindromic motifs are present upstream of *hrtBA_Ef_*. The 2 palindromes are in blue (P1) and in red (P2); the RBS sequence is underlined and the start codon is in green. **(B)** FhtR binds to the promoter region of *hrtBA_Ef_*. EMSA shows binding of FhtR to P_hrtBA_. 0.25 pmoles of the *hrtBA_Ef_* promoter fragment were incubated with increasing amounts of MBP-FhtR as indicated in molar ratios. DNA shift was visualized with GelRed (Biotium) following PAGE. C1, C2 indicate the 2 shifted DNA-protein complexes and NC, the noncomplexed DNA. Result is representative of at least 3 independent experiments. **(C)** Role of the 2 palindromic sequences on FhtR binding to the promoter region of *hrtBA_Ef_*. EMSA was performed as in (B) with either the native or P_hrtBA_ DNA fragment (as in A) or mutated versions fragments P_hrtBA P1*, P2*_, P_hrtBA P1*_, P_hrtBA P2*_ containing P1 and P2 randomly exchanged nt P1* and P2* respectively (A). MBP-FhtR: DNA (2.5 pmol: 0.25 pmol respectively). Result is representative of at least 3 independent experiments. **(D)** Effect of hemin on binding of FhtR to the *hrtBA_Ef_* promoter. 0.25 pmoles of the *hrtBA_Ef_* promoter DNA was incubated with 2.5 pmoles of MBP-FhtR (MBP-FhtR: DNA) together with increasing amounts of hemin as indicated (molar ratios) and analyzed by EMSA as in (B). Result is representative of at least 3 independent experiments. **(E**) Substitution of the 2 palindromic nt sequences, P1 and P2, in P_hrtBA_ abrogates FhtR mediated control of *hrtBA_Ef_* transcription. WT OG1RF strain was transformed either with the reporter plasmid pP_hrtBA_-*lac* or pP_hrtBA P1*P2*_ *-lac* with P1 and P2 randomly substituted nt as in (B). B-gal activity was determined as in Fig. 1 following incubation with 2.5 μM hemin. Results represents the average ± S.D. from triplicate technical samples. ****, *P* < 0.0001, Student’s *t* test; ns, non significant, Student’s *t* test. Result is representative of 3 independent experiments. (**F**) Substitution of either P1 or P2 nt sequences in P_hrtBA_ enhances its transcriptional activation by hemin. WT OG1RF strain was transformed either with the reporter plasmid pP_hrtBA_-*lac* or pP_hrtBA P1*_ *-lac* or pP_hrtBA P2*_ *-lac* with P1 and P2 randomly substituted nt as in (B). B-gal activity was determined as in Fig. 1 following incubation with 0.1 and 0.5 μM hemin. Results represent the average ± S.D. from triplicate technical samples. ****, *P* < 0.0001; ***, P=0.0003; **, P = 0.0028; ns, non significant, one-way ANOVA followed by Tukey’s multiple comparisons test. Result is representative of 3 independent experiments.

We then tested the effects of heme on FhtR binding by EMSA. Addition of hemin to MBP-FhtR abolished the formation of DNA-FhtR complex, as seen by the progressive disappearance of band shifts with increasing hemin concentrations (Fig. 5D). Complete release of FhtR from P_hrtBA_ was obtained when hemin was in 10-fold molar excess over FhtR. Both C1 and C2 complexes were revealed when intermediate amounts of heme were added (0.1 and 1 μM; Fig. 5D). This suggests the release of MBP-FhtR from only one operator depending on the saturation level of FhtR with hemin.

The role of each operator in the control of *hrtBA_Ef_* was investigated *in vivo*, using P_hrtBA_ or a P_hrtBA P1*, P2*_ promoter variant to control the *lac* gene (Fig. 5E). In contrast to pP_hrtBA_-*lac*, which was strongly induced with 1 μM hemin, P_hrtBA P1*, P2*_-*lac* was constitutively expressed (Fig. 5E). In the absence of heme, both P1 and P2 were required for full P_hrtBA_ repression by FhtR (Fig. 5F). We propose that the presence of 2 operators provides strong repression of the *hrtBA_Ef_* promoter, which prevents transcriptional leakage and allows for fine tuning of HrtBA_*Ef*_ expression. Taken together, these results demonstrate that FhtR is a heme sensor that directly controls heme homeostasis by regulating *hrtBA_Ef_* transcription.

### FhtR controls HrtBA_*Ef*_, the gatekeeper of intracellular heme availability

Our observation that FhtR regulates intracellular heme pools even at low heme concentrations, led us to hypothesize that FhtR controls intracellular heme availability in *E. faecalis*. We tested this possible role of FhtR on the *E. faecalis* endogenous heme-dependent catalase (KatA). While *katA* transcription is not susceptible to heme induction, KatA protein stability relies on the presence of heme (9, 11). KatA mediated H_2_O_2_ catalysis was measured in WT, Δ*fhtR* and Δ*fhtR*(p*fhtR*) strains (Fig. 6A). In the absence of hemin, H_2_O_2_ consumption was at a basal level (Fig. S4A), thus excluding major contributions of other enzymes in our conditions. In the presence of 1 μM hemin (Fig. 6A), the Δ*fhtR* mutant exhibited about 30 % catalase activity compared to WT and complemented Δ*fhtR*(p*fhtR*) strains, as evaluated by the percent of catabolized H_2_O_2_ (Fig. 6A). This was further confirmed by comparing amounts of KatA (holoKatA) by WB, using anti-KatA antibody (kindly provided by L. Hederstedt). In the absence of hemin, KatA was expressed at low levels in WT, Δ*fhtR* and Δ*fhtR*(p*fhtR*) strains (Fig.6B). Comparatively, addition of hemin strongly increased the amounts of KatA in WT and complemented Δ*fhtR*(p*fhtR*) strains, but not in the Δ*fhtR* mutant (Fig. 6B). Low KatA availability in Δ*fhtR* is readily explained by constitutive heme efflux (*via* HrtBA_*Ef*_), and consequently depleted intracellular heme pools in this mutant. We then evaluated the survival capacity of *E. faecalis* OG1RF WT, Δ*fhtR* and Δ*fhtR*(p*fhtR*) strains when challenged with 2.5 mM H_2_O_2_. In the absence of hemin, all strains grew equivalently without H_2_O_2_ (Fig. S4B). In contrast, while hemin addition rescued the survival of both WT and Δ*fhtR*(p*fhtR*), the Δ*fhtR* strain remained hypersensitive to H_2_O_2_. Deletion of *hrtBA_Ef_* in the Δ*fhtR* strain (Δ*fhtR*Δ*hrtBA_Ef_*) restored the survival capacity in presence of hemin (Fig. S4C). Thus, poor survival of Δ*fhtR* reflects the lack of heme needed to stabilize KatA (Fig. 6D). Finally, the OG1RF mutant (*katA::tetR*) was hypersensitive to hemin toxicity showing that KatA was also responsible for mediating oxidative stress generated by hemin (Fig. 4E). Taken together, these results identify FhtR as the direct and indirect regulator of HrtBA_*Ef*_-mediated heme-efflux and KatA activity respectively; both mechanisms lowering heme stress in *E. faecalis* OG1RF. They show that FhtR is a key mediator of heme homeostasis, and consequently, of oxidative stress response in *E. faecalis* generated by H_2_O_2_.

**Figure 6.**
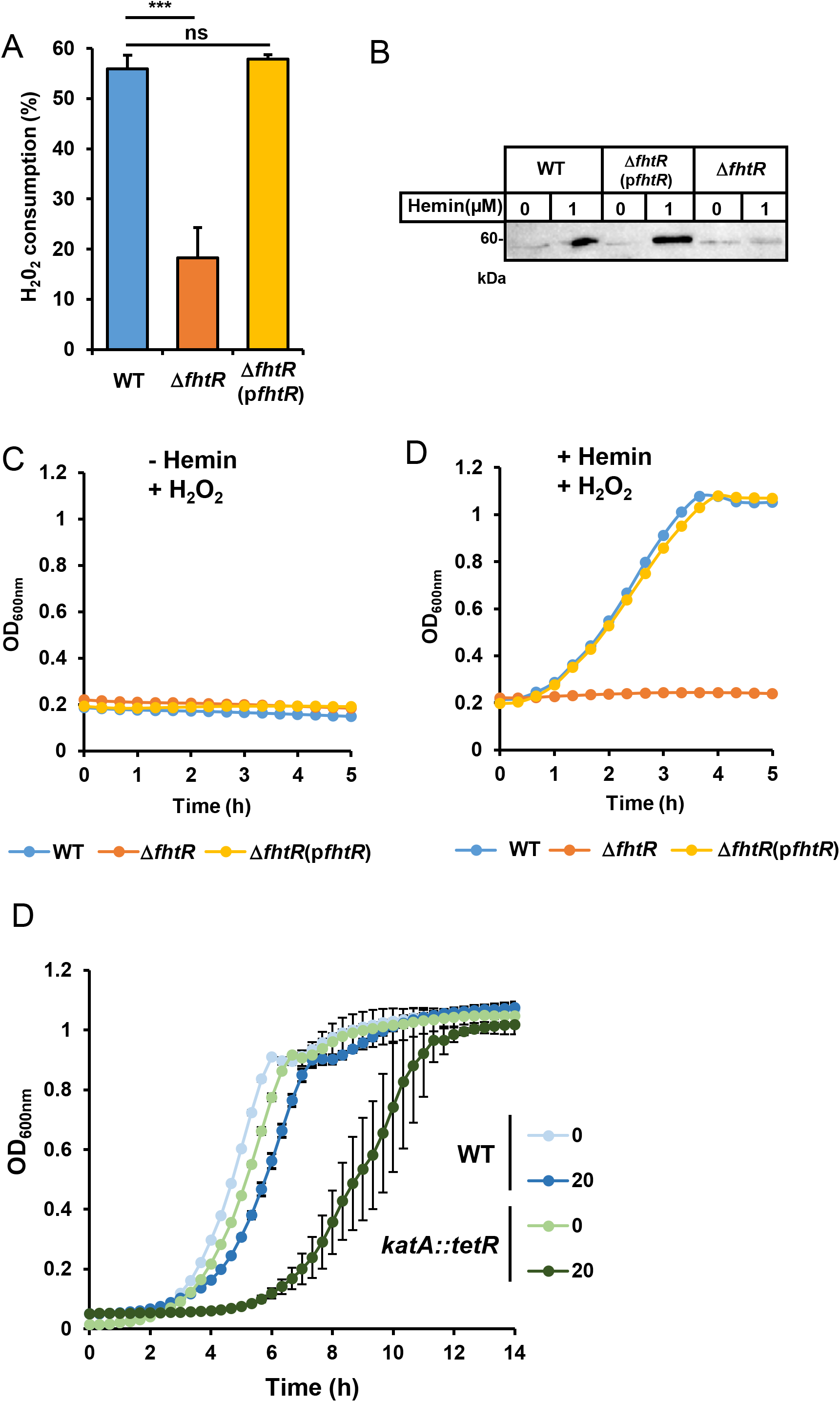
FhtR controls intracellular utilization of heme. **(A) *fhtR* deletion limits heme-dependent KatA activity**. KatA enzymatic activity in *E. faecalis* was assessed on WT, Δ*fhtR* and Δ*fhtR*(p*fhtR*) grown ON with 1 μM hemin. Catalase activity was determined on equivalent number of bacteria incubated with 100 μM H_2_O_2_ for 30 min with the spectrophotometric FOX1 method based on ferrous oxidation in xylenol orange. Results expressed as the % of H_2_O_2_ metabolized in respective strains grown without hemin and are the average ± S.D. of technical triplicates. ***, *P* = 0.0001; ns, non significant, one-way ANOVA followed by Dunnett’s posttest comparison to WT. Result is representative of 3 independent experiments. (**B**) Expression of KatA is impaired in the Δ*fhtR* OG1RF mutant. KatA was followed on immunoblots with a polyclonal anti-KatA antibody on equivalent amounts of protein (20 μg) from lysates of WT, Δ*fhtR* and Δ*fhtR*(p*fhtR*) strains as in (A) were separated on SDS-PAGE and probed with an anti-KatA polyclonal antibody. The presented results are representative of 3 independent experiments. **(C, D)** Δ*fhtR* mutant is hypersensitive to H_2_O_2_. ON cultures of WT, Δ*fhtR* and Δ*fhtR*(p*fhtR*) strains were diluted to OD_600nm_=0.01 and grown to OD_600nm_=0.5 in the absence (C) or presence of 1 μM hemin (D). Cultures were distributed in a 96 well plate and supplemented with 2.5 mM H_2_O_2_ at 37 °C in a microplate Spark spectrophotometer (Tecan). OD_600nm_ was measured every 20 min. Results represent the average ± S.D. from triplicate technical samples and is representative of 3 independent experiments. (**E**) KatA limits heme toxicity. ON cultures of WT and *katA::tetR* OG1RF strains were diluted to OD_600nm_=0.01 and grown in 96 well plate in presence of the indicated concentration of hemin. Results represent the average ± S.D. from triplicate technical samples and is representative of 3 independent experiments.

### Heme sensing in the gastrointestinal tract

*Enterococcus faecalis* is a normal resident of the gastrointestinal tract (GIT) of vertebrates, an ecosystem where heme is available (27–30). We therefore investigated whether *hrtBA_Ef_*-mediated heme management is required by *E. faecalis* in the GIT in a murine gastrointestinal model. We generated *E. faecalis* OG1RF strains expressing the *luxABCDE* (*lux*) operon from *Photorhabdus luminescens* driven by: i-P_hrtBA_ (pP_hrtBA_-*lux*), which emits light specifically in the presence of hemin (Fig. 7A), ii-a constitutive promoter P23 (p*lux*), constitutively emitting light for bacterial tracking (13), or iii-a control promoterless vector, pP_∅_-*lux*. Cultures of these strains were orally inoculated in the digestive tract of mice, and light emission from whole live animals was measured in an *in vivo* imaging system IVIS200, 6 h post-inoculation (Fig. 7B). This time delay corresponded to the maximum light emission from the tracking strain OG1RF(p*lux*) (Fig. S5A). Luminescence signaling from the ingested *E. faecalis* pP_hrtBA_-*lux* heme sensor strain also localized in the abdomen, similarly to the tracking strain (Fig. 7B). Examination of dissected organs revealed that the heme sensor-associated luminescence was mainly detected in the caecum (Fig. 7C), correlating with the high bacterial load of this organ (WT(p*lux*), Fig. 7C). A significant signal was also detected in the feces from inoculated animals further highlighting that *E. faecalis* was able to scavenge and internalize heme within the digestive tract to induce *hrtBA_Ef_* expression (Fig. 7D). Finally, mice as well as human feces samples (as well as faecal waters (Fig. S6A and S6B)) from healthy individuals were able to induce luminescence from WT(pP_hrtBA_-*lux*) *in vitro* excluding the possibility that induction of P*_hrtBA_ in vivo* could result from *E. faecalis* inoculation to the animals (Fig. 7E). Therefore, FhtR heme sensor activity is active and relevant to *E. faecalis* heme management in the lumen of the GIT.

**Figure 7.**
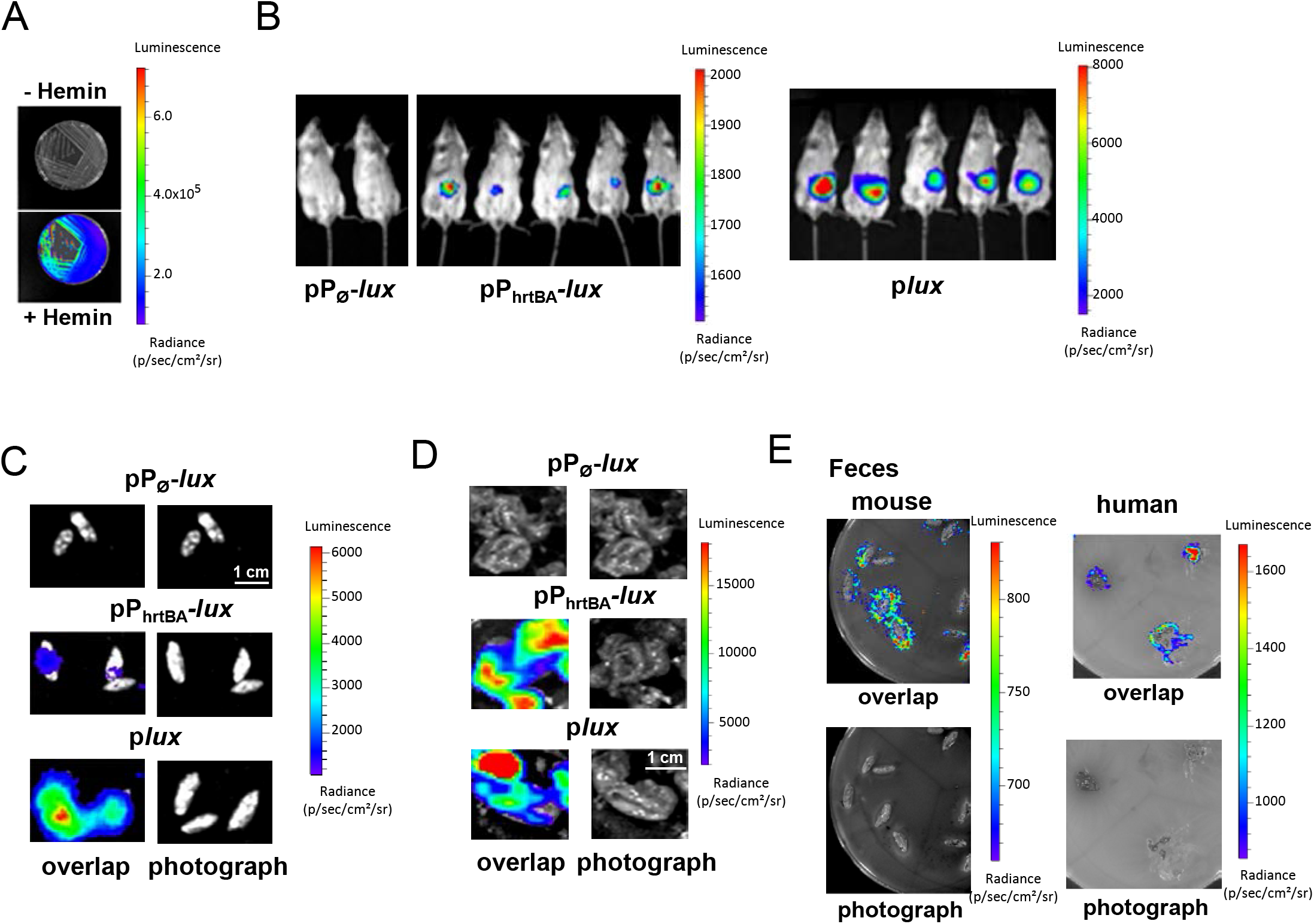
HrtBA_*Ef*_ is induced in the gastrointestinal environment. **(A)** Heme dependent light emission by the heme sensing OG1RF strain. OG1RF WT(pP_hrtBA_-*lux*) was on M17G agar plates ± 20 μM hemin. Plates were incubated at 37 °C for 24 h and luminescence was visualized using an IVIS 200 luminescence imaging system, (acquisition time, 1 min; binning 8). **(B)** Heme sensing by *E. faecalis* over the course of infection. Female BALB/c mice were force-feeded with 10^8^ CFU of WT(pP_∅_-*lux*) (control strain), WT(pP_hrtBA_-*lux*) (sensor strain) or WT(pl*ux*)(tracking strain). At 6 h post-inoculation, anesthetized mice were imaged in the IVIS 200 system (acquisition time, 20 min; binning 16). Figure shows representative animals correponding to a total of 15 animals for each condition in 3 independant experiments. (**C**) Caecums exhibit high heme sensing signal. Animals as in (B) were euthanized and immediately dissected. Isolated GIT were imaged as in (B). Caecums that exhibited most of the luminescence are shown (acquisition time, 5 min; binning 8). Scale= 1 μM. (**D**) Visualization of heme sensing in feces collected from mice following ingestion of WT(pP_∅_-*lux*), WT(pP_hrtBA_-*lux*) or WT(p*lux*). WT(pP_hrtBA_-*lux*) as in (B) were collected after 6-9h following gavage. Feces were imaged as in (B) (acquisition time, 20 min; binning 16). Scale = 1 μM. (**E**) Human and mice feces samples activate heme sensing. Human feces from 3 healthy human laboratory volunteers and mice feces from 6 months old female BALB/c mice were deposited on M17G agar plates layered with soft agar containing WT OG1RF (pP_hrtBA_-*lux*). Plates were incubated at 37 °C for 16 h and imaged in the IVIS 200 system (acquisition time, 10 min; binning 8). Figure shows representative results corresponding to a total of 3 independent experiments.

### Heme sources for *E. faecalis* in the GIT

The results described above implies that *E. faecalis* internalizes heme in the intestinal environment to activate FhtR. Thus, an interesting question remains as to identify heme sources that are accessible to *E. faecalis* in the GIT. Normal bleeding (occult blood), exacerbated in intestinal pathologies, as well as food (as meat) are considered as main sources of heme within the GIT (the second being excluded in mice) (27–30). We thus tested if hemoglobin (Hb) or blood could induce P_hrtBA_ from WT OG1RF(pP_hrtBA_-*lux*) (Fig. 8A). Luminescence from heme sensor plasmid was induced in *E. faecalis* at the proximity of Hb and blood deposits similarly to hemin (Fig. 8A) suggesting that heme associated with physiologically available heme sources are accessible to internalization by *E. faecalis*. Cross-feeding of metabolites, inclusing heme between bacteria has also been reported (27, 31). The possibility that *E. faecalis* could scavenge heme from heme synthesizing bacteria, such as *E. coli*, a phylum that becomes prevalent together with *E. faecalis* throughout dysbiosis, was evaluated by growing WT OG1RF carrying pP_hrtBA_-*lux* heme sensor plasmid with *E. coli*. (Fig. 8B). Strikingly, induction of luminescence was localized to areas of overlap between the two bacteria suggesting that *E. coli* was a heme donor of heme cross-feeding between the 2 bacteria when in close contact. Thus, heme crossfeeding between bacterial symbionts in the gut might provide a heme source for *E. faecalis*. We conclude that *E. faecalis* heme sensor is activated by the heme sources available in the GIT.

**Figure 8.**
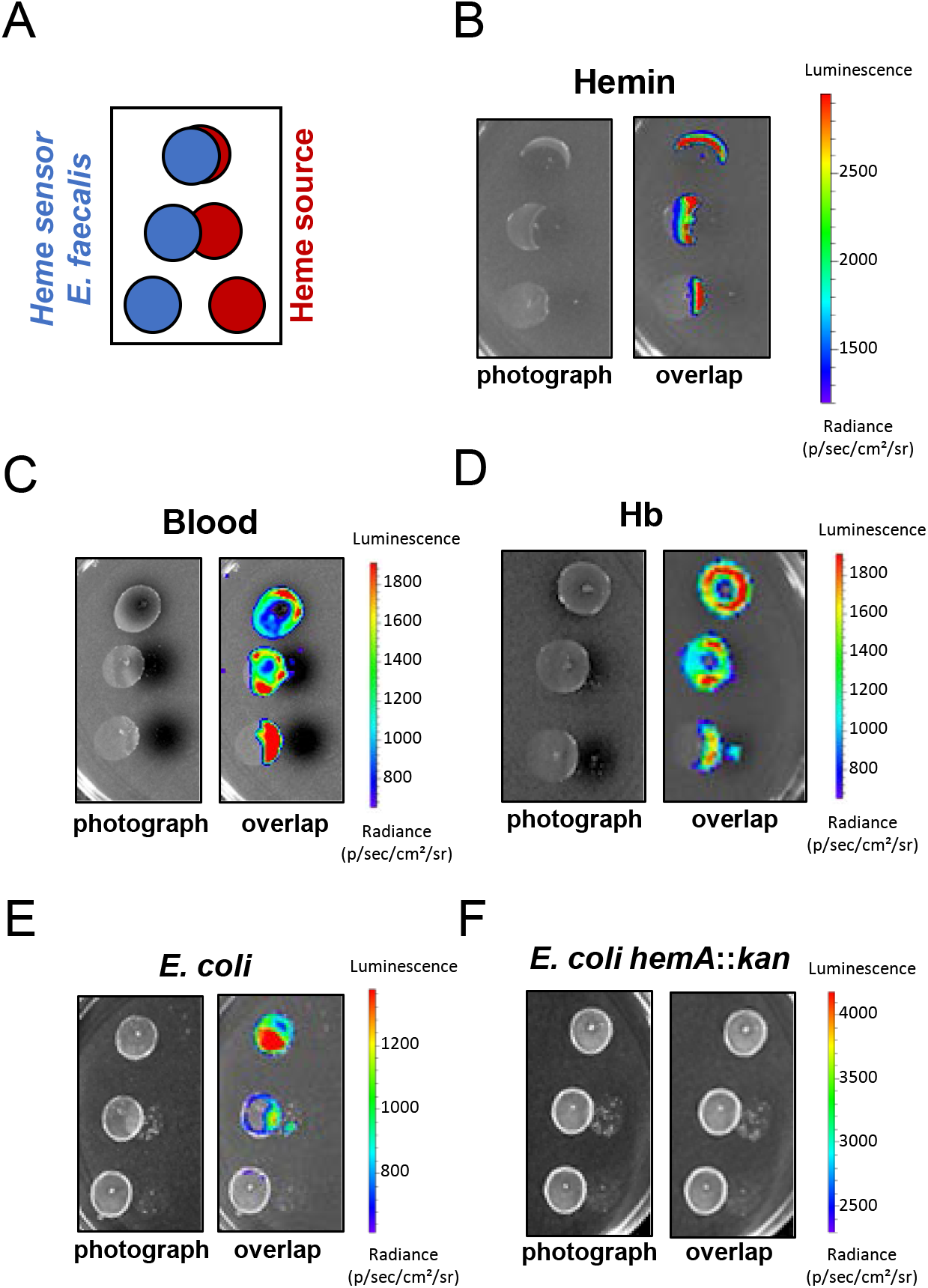
Heme sources for *E. faecalis* in the GIT. **(A)** Intracellular heme sensing setup. 10 μl deposits of the indicated heme sources were realized on M17G solid medium as shown in red. WT OG1RF carrying pP_hrtBA_-*lux* heme sensor plasmid was plated as 10 μl spots at OD_600nm_=0.01 as shown in blue. Plates were incubated at 37 °C for 16 h and imaged in the IVIS 200 system (acquisition time, 10 min; binning 8). Figures [B-F] show representative results corresponding to a total of 3 independent experiments. **(B)** Visualization of heme sensing from hemin deposits. 1 mM hemin in PBS was used. (**C, E**) Heme from blood (bovine), from hemoglobin (human) are heme sources for *E. faecalis*. Heparanized bovine blood (Thermofisher) and freshly dissolved human hemoglobin (1 mM) in PBS were used. (**F, G**) *E. coli* is a heme donor for *E. faecalis. E. coli* (NEB10, New England Biolabs) or a mutant strain that cannot synthesize heme (*hemA::kan*) mutant at OD_600nm_=0.1 were platted as in (A). Only the heme synthesizing strain was able to cross-feed heme to *E. faecalis*.

## Discussion

*E. faecalis* is a core member of the microbiome, and is also the cause of a variety of severe infections (32). The central role of heme in reprogramming *E. faecalis* metabolism and fitness led us to investigate how heme homeostasis is controlled. A novel heme sensor, FhtR, is shown here to regulate heme intracellular homeostasis in *E. faecalis*. FhtR-heme complexes de-repress the *hrtBA_Ef_* operon, leading to the HrtBA-mediated management of intracellular heme pools. While expression of HrtBA is a conserved strategy in multiple Gram-positive organisms, *E. faecalis* appears to be the first example of an opportunistic pathogen where HrtBA is not controlled by the two-component system HssRS. Blast analysis of FhtR homologs in several Gram-positive bacteria showed that the regulator is present only in enterococci, vagococci and carnobacteria (Fig. S7A and Fig. S7B for FhtR alignments and phylogenic tree). FhtR shares no homology with HrtR, a TetR regulator of *hrtRBA* in *Lactococcus lactis* (18). In contrast to HrtR which autoregulates its own expression, *fhtR* is monocistronic and expressed constitutively, implying that only HrtBA_*Ef*_ expression is controlled by heme (18).

We characterized FhtR as a heme-binding protein through pentacoordinated ligation of the heme iron implying a tyrosine. This state of coordination is mostly found in heme receptors that transiently bind heme, such as IsdA, IsdC, and IsdH in *S. aureus*, or HmA in *Escherichia coli* (33). FhtR blocks *hrtBA_Ef_* transcription by binding to two distinct 14-nt inverted repeats sequences in its promoter region. Alleviation of repression occurs when the heme-FhtR complex loses its affinity for its DNA binding sites. Conformational changes upon ligand binding is a shared mechanism among TetR regulators, leading to uncoupling from DNA (24). We thus hypothesize that these events, which we verified *in vitro*, explain FhtR regulation of the *hrtBA_Ef_* efflux pump in *E. faecalis*. The unique features of FhtR in *E. faecalis* compared to other regulators of *hrtBA* genes encoding efflux pumps support the idea that control of HrtBA-mediated heme homeostasis may vary among bacteria as a function of their lifestyle. It is thus tempting to speculate that differences in host niches, and in heme utilization and metabolism, might explain disparities in bacterial heme sensing mechanisms.

Heme efflux by HrtBA is reported as a bacterial detoxification mechanism that prevents intracellular heme overload (8, 13, 15, 16, 18). We showed here that HrtBA induction is required for *E. faecalis* survival when heme concentrations reached toxic levels (> 25 μM). Yet, *hrtBA_Ef_* was induced at heme concentrations as low as 0.1 μM, suggesting that heme efflux is also needed at non-toxic levels. Interestingly, *E. faecalis* encodes (at least) two heme-dependent enzymes, cytochrome *bd* oxidase and catalase (9, 10, 34). Both these enzymes not only bind heme and thus lower free heme levels, but also actively lower oxidative stress generated by heme. It will be of interest to determine the hierarchy of heme binding between FhtR, cytochrome *bd* oxidase, and catalase *in vivo.*

To date, no heme import function has been identified in *E. faecalis*, nor in other tested Gram-positive heme auxotrophs (13, 18). Blast analysis of these bacteria failed to identify genes of the *isd* heme import system described in *Staphylococcus aureus* (35, 36). In *S. aureus*, heme receptors and transporters are induced in iron-depleted growth media, and imported heme is used as an iron source (36). Thus, our findings led us to question the need for a dedicated transport system to internalize exogenous heme in *E. faecalis*, and to propose an alternative hypothesis. We noted that HrtBA_*Ef*_ is a member of the MacB family of efflux-pumps that is distinct from other structurally characterized ABC transporters (37). A model based on MacB transport of antibiotics and antimicrobial peptides in *Streptococcus pneumoniae* proposed that transmembrane conformational changes promote lateral entry of substrates in the membrane before they reach the cytoplasm (38). Based on the previous and present data (22), we propose that HrtB_*Ef*_ has the integral role as heme “gatekeeper” in the cell. Like MacB antibiotics and antimicrobial substrates (39), membrane-bound heme could either enter passively into the intracellular compartment, and or be effluxed by HrtB before this step. Altogether, our results place HrtBA_*Ef*_ at the forefront of heme acquisition in *E. faecalis. E. faecalis* is thus dependent on the key role of FhtR to adapt to the dichotomy between toxicity and benefits of heme which may be crucial in the host.

*In vivo* bioluminescent imaging of *E. faecalis* using an FhtR-based sensor identified the GIT as an environment where HrtBA_*Ef*_ is expressed. The gut lumen of healthy individuals contains heme, independently of the nature of ingested food or of the microbiota (27–30). Heme in the GIT is reported to mainly originate from Hb from normal bleeding (occult blood) (40). Accordingly, *E. faecalis* was able to internalize heme from blood and Hb *in vitro*. In addition, a common microbiota constituent, *Escherichia coli*, is shown to be a heme donor, suggesting a novel basis for intestinal bacterial interactions. As several phylae composing the core microbiota are heme-auxotrophs with vital heme requirements, it is tempting to hypothesize that normal or disease associated fluctuations in host heme levels could be detected by FhtR to adjust its intracellular level and optimize bacterial fitness. Interestingly, *E. faecalis* causes a variety of severe infections, most often among antibiotic-treated hospitalized patients with intestinal dysbiosis favorizing high *E. coli* and enterobacterial populations (41). Taken together our results suggest that the FhtR-sensor and the HrtBA_*Ef*_ heme gatekeeper allow *E. faecalis* to optimize its adaptation to variable heme pools in the host.

## Materials and Methods

### Bacterial strains and growth conditions

Strains and plasmids used in this work are listed in Table S1. *E. coli* NEB 10 (New England Biolabs) was grown in LB medium and *E. coli* C600 *hemA*::kan in M17 medium supplemented with 0.5 % glucose (M17G). Experiments in *E. faecalis* were all performed using strain OG1RF and derivatives (Table S1). *E. faecalis* was grown in static conditions at 37°C in M17G. When needed, antibiotics were used for *E. coli* at 50 μg.ml^−1^ kanamycin and 100 μg.ml^−1^ ampicilin, respectively; for *E. faecalis* at 30 μg.ml^−1^ erythromycin. Oligonucleotides used for plasmid constructions are listed in Table S2. Hemin (Fe-PPIX) (Frontier Scientific) was prepared from a stock solution of 10 mM hemin chloride in 50 mM NaOH. In this report, heme refers to iron protoporphyrin IX regardless of the iron redox state, whereas hemin refers to ferric iron protoporphyrin IX. For growth homogeneity, WT and mutants strains were transformed with the promoterless pTCV-*lac* plasmid when compared to complemented strains. Plasmid construction and *E.faecalis* gene deletion are described in Supplemental Material and Methods (Text S1).

### β–galactosidase assays

Stationary phase cultures were diluted at OD_600nm_ = 0.01 in M17G and grown to OD_600nm_ = 0.5. Hemin was added to cultures, further grown for 1.5 h. β–galactosidase activity was quantified by luminescence in a Spark microplate luminometer (TECAN, Austria) using the β-glo substrate (Promega) as described (18)

### Bacterial lysate preparation

Bacteria were pelleted at 3,500g for 10 min, resuspended in 20 mM Hepes pH 7.5, 300 mM NaCl, and disrupted with glass beads (Fastprep, MP Biomedicals). Cell debris were removed by centrifugation at 18,000 g at 4 °C for 15 min from the bacterial lysate supernatant. Proteins were quantified with the Lowry Method (Biorad).

### Heme concentration determination in bacterial lysates

Equivalent amounts of proteins (in a volume of 250 μl) were mixed with 250 μl of 0.2 M NaOH, 40 % (v/v) pyridine and 500 μM potassium ferricyanide or 5 μl of 0.5 M sodium dithionite (diluted in 0.5 M NaOH) and 500-600 nm absorption spectra were recorded in a UV-visible spectrophotometer Libra S22 (Biochrom). Dithionite-reduced minus ferricyanide-oxidized spectra of pyridine hemochromes were used to determine the amount of heme *b* by following the value of the difference between absorbance at 557 nm and 540 nm using a difference extinction coefficient of 23.98 mM^−1^cm^−1^ (42).

### Recombinant MBP-FhtR purification

*Escherichia coli* Top10 or C600 Δ*hemA* mutant strain carrying pMBP-FhtR or pMBP-FhtR^Y132F^ was used for the recombinant production of the fusion proteins (Table S2). The protein was purified by affinity chromatography on amylose resin as reported previously (18). Briefly, *E. coli* Top10 or C600 Δ*hemA* strains were grown to OD600 = 0.6 or 0.3 respectively and expression was induced with 1 mM isopropyl 1-thio-β-D-galactopyranoside (IPTG) ON at RT. Cells were pelleted at 3500 g for 10 min, resuspended in 20 mM Hepes pH 7.5, 300 mM NaCl, containing 1 mM EDTA (binding buffer), and disrupted with glass beads (Fastprep, MP Biomedicals). Cell debris was removed by centrifugation at 18,000 g for 15 min at 4°C. MBP-tagged proteins were purified by amylose affinity chromatography (New England Biolabs) following manufacturer’s recommendations: the soluble fraction was mixed with amylose resin and incubated on a spinning wheel at 4°C for 1 h. The resin was then centrifuged, and washed 3 times with binding buffer. Purified proteins were eluted in binding buffer containing 10 mM maltose, and dialyzed against 20 mM Hepes pH 7.5, 300 mM NaCl.

### Heme-dependent catalase expression and activity

KatA expression was followed on immunoblots with a polyclonal anti-KatA antibody (9). Catalase activity was determined on whole bacteria incubated with 100 μM H_2_O_2_ with the spectrophotometric FOX1 method based on ferrous oxidation in xylenol orange as described previously (43, Wolff, 1994 #22).Absorption was measured at 560 nm.

### Electrophoretic Mobility Shift Assay (EMSA)

A 325-bp DNA fragment containing the *hrtBA_Ef_* promoter (P_hrtBA_) was amplified by PCR from *E. faecalis* OG1RF genomic DNA with primers pair (O21-O22) (Table S2). In P_hrtBA_, the two 14-nt palindromic sequences; P1: 5’-TTATCAATCGATAA-3’ and P2: 5’-TTATCGATTGATAA-3’ were randomly altered to P1*: 5’-ACTTGTATACATAA-3’ and P2*: 5’-ATATCTTGTATAAG-3’ to generate 3 DNA variants P_hrtBA P1*_, P_hrtBA P2*_ and P_hrtBA P1*, P2*_, cloned into pUC plasmid (pUC-VS1, pUC-VS2 and pUC-VS3, Table S1) (Proteogenix, France) that were used as templates to PCR amplify the promoter region DNA variants with the primer pairs (O21-O22) (Table S2). A DNA fragment located in the *hrtBA_Ef_*-coding region was amplified with (O23-O24) primers (Table S2) and used as a negative control. Interaction between binding studies were performed in 20 mM Tris-HCl, pH 8, 50mM KCl, 0.2 mM MgCl_2_, 1 mM EDTA, 0.2 mM DTT, and 5% glycerol. Reaction mixtures were incubated for 1 h at 37 °C as reported previously (18). Binding was analyzed by gel electrophoresis on a 7 % polyacrylamide gel in TBE buffer stained with GelRed (Biotium) following electrophoresis.

### Ethics statement

Animal experiments were carried out in strict accordance with the recommendations in the guidelines of the Code for Methods and Welfare Considerations in Behavioural Research with Animals of the EEC council (Directive 2010/63/EU). The protocols were approved by the Animal Care and Use Committee at the Research center of Jouy en Josas (COMETHEA; protocol number 15–61) and by the Ministry of Education and Research (APAFIS#2277-2015081917023093 v4). All efforts were undertaken to minimize animal suffering. All experimental procedures were performed in biosafety level 2 facilities.

### *In vivo* heme sensing assay in the mouse GIT

For inoculation in the digestive tract, *E. faecalis* strains were prepared as follows: OG1RF precultures were diluted and grown in M17G to OD_600nm_ = 0.5 that was determined to correspond to 6.10^8^ CFU/ml. Bacteria were then centrifuged at 6000 rpm at 4°C for 15 min and pellets were resuspended in PBS to a final concentration of 2.10^8^ cells/ml. Bacterial stocks were aliquoted and frozen in liquid nitrogen. Aliquots were kept at −80 °C until use. Bacterial counts were confirmed by plating serial dilutions of cultures. 6 week old female BALB/c mice (Janvier, France) were orally administered by gavage of 10^8^ using a feeding tube. Image acquisition of isoflurane anesthesized mice was performed at the indicated time following gavage. Following image acquisition, mice were removed from the IVIS 200 imaging system and immediately sacrificed by cervical dislocation. When indicated, the animals were dissected for imaging of the isolated organs. *In vivo* luminescence imaging procedure are described in Supplemental Material and Methods (Text S1).

## Acknowledgments

This work was supported by the HemeDetox - 17-CE11-0044-01 project by the French “Agence Nationale de la Recherche”. V. Saillant is the recipient of a doctoral fellowship from the French ministry of Research and Paris-Saclay University. The funders had no role in study design, data collection and analysis, decision to publish, or preparation of the manuscript. We thank Dr. A. Gruss (INRAE, France) and Dr. P. Moenne-Loccoz, Oregon Health and Science University, USA) for their help and valuable comments on this manuscript. We thank Dr. M. Vos, Dr. U. Liebl (CNRS, INSERM, Ecole Polytechnique, France) and Dr. P. Delepelaire (IBPC, France) for their technical help and insightful comments on our work. We are grateful to Dr. Lars Hederstedt (Lund University, Sweden) for the generous gift of the anti-KatA antibody and OG1RF mutants.

## Abbreviations

Bgal: B-galactosidase
ECL: Enhanced chemiluminescence
EMSA: electromobility shift assay
EPR: electron paramagnetic resonance
FhtR: Faecalis heme transport regulator
GIT: gastrointestinal tract
Hb: hemoglobin
HrtA: heme regulated transport A
HrtB: heme regulated transport B
HrtR: heme regulated transport R
IPTG: isopropyl β-D-1-thiogalactopyranoside
KatA: catalase A
PAGE: polyacrylamide gel electrophoresis
Xgal: 5-bromo-4-chloro-3-indolyl-beta-D-galactopyranoside
WB: Western blot

## Supplemental material

### Supplemental materials and methods

#### Plasmid construction

The plasmids [1–9] were obtained with similar cloning strategy: PCR amplification of the described DNA inserts with oligonucleotides from Table S2, digestion with (*Eco*RI, *Bam*HI) and ligation with the (*Eco*RI, *Bam*HI) digested plasmid pTCV-*lac* (1) (Table S1). For the following plasmids: [1], pP_hrtBA_-*lac* (pTCV-VS2); [2], pP_fhtR_-*fhtR*, P_hrtBA_-*lac* (pTCV-VS3) and [3], pP_fhtR_-*lac* (pTCV-VS4); fragments containing either P_hrtBA_, P_fhtR_-*fhtR*-P_hrtBA_ or P_fhtR_ DNA sequences were PCR amplified with the primers pairs (O1-O2), (O2-O3) or (O3-O4) respectively. Plasmid [4], pP_fhtR_-*fhtR*-HA, P_hrtBA_-*lac* (pTCV-VS5) was obtained by PCR amplification of 2 DNA fragments, P_fhtR_-*fhtR* and P_hrtBA_ using primer pairs (O3-O5) and (O2-O6), respectively, followed by their fusion by PCR-overlap with the primers (O2-O3) leading to the addition of nt encoding the hemagglutinin influenza epitope (HA, YPYDVPDYA) at the 3’ end of *fhtR* (2). The mutation Y132F was introduced into the coding sequence of *fhtR* in the plasmids [5], pP_fhtR_-*fhtR*^Y132F^ (pTCV-VS6) and [6], pP_fhtR_-*fhtR*^Y132F^-HA (pTCV-VS7) by a PCR-overlap with (O2-O3) of 2 PCR products obtained with primer pairs (O3-O7) and (O2-O8) using pTCV-VS3 and pTCV-VS5 as templates, respectively. Plasmids [7], pP_hrtBA P1*_-*lac* (pTCV-VS8); [8], pP_hrtBA P2*_-*lac* (pTCV-VS9) and [9], pP_hrtBA P1*P2*_-*lac* (pTCV-VS10) were obtained by PCR amplification of P_hrtBA_ nt sequence with primers pair (O1-O2) using pUC-VS1, pUC-VS2 and pUC-VS3 as templates, respectively (Table S1). The plasmid pP_hrtBA_-*lux* (pTCV-VS11) was constructed as described for pP_hrtBA_-*lac* using the plasmid vector pTCV-J22 (Table S1). Plasmids pMBP-FhtR and pMBP-FhtR^Y132F^ (Table S1) were obtained by PCR amplification of FhtR ORF from *E. faecalis* OG1RF genomic DNA and pP_fhtR_-*fhtR*^Y132F^ respectively, with the primers pairs (O9-O10) (Table S2). The resulting fragments were digested with *Eco*RI and *Pst*I and ligated with the plasmid pMAL-c4X (New England Biolabs) (Table S1). The 3 plasmids pΔ*hrtBA* (pG1-VS1), pΔ*fhtR* (pG1-VS2) and pΔ*fhtR*Δ*hrtBA* (pG1-VS3) were constructed by PCR amplification of 2 fragments of ~ 600-750 pb flanking the genes OG1RF_RS02770-OG1RF_RS02775 (*hrtBA*); OG1RF_RS02765 (*fhtR*) gene and OG1RF_RS02765-OG1RF_RS02775 (*fhtR*-*hrtBA*) with the respective (O11-O12) and (O13-O14); (O15-O16) and (O17-O18); (O15-O19) and (O14-O20) oligonucleotides (Table S1 and S2). Each pair of fragments was fused by PCR overlap as above with the DNA primers pairs (O11-O13); (O15-O18) and (O14-O15), respectively (Table S2). The resulting fragments were digested with *Bam*HI and *Hind*III and ligated into the thermosensitive pG1 (pG1-VS2, Table S1). All plasmids were verified by DNA sequencing.

#### *E. faecalis* Δ*hrtBA*, Δ*fhtR* and Δ*fhtR*Δ*hrtBA* mutants

The plasmids pΔ*hrtBA* (pG1-VS1), pΔ*fhtR* (pG1-VS2) and pΔ*fhtR*Δ*hrtBA* (pG1-VS3) (Table S1) were transformed by electroporation in *E. faecalis* OG1RF strain. The double cross-over events leading to Δ*hrtBA*, Δ*fhtR and* mutants and Δ*fhtR*Δ*hrtBA* double mutant were obtained as described (3). Correct inactivation of the targeted genes was confirmed by sequencing.

#### Titration of MBP-FhtR with hemin

ApoFhtR/heminstochiometry: hemin binding affinities of MBP-FhtRwere was determined by adding 0.5 to 1 μl increments of a 200 μM hemin solution to cuvettes containing 100 μl test samples of 20 μM MBP-FhtR in 20 mM Hepes pH 7.5, 300 mM NaCl, or reference sample without protein. Spectra were measured from 350 nm to 700 nm in a UV-visible spectrophotometer Libra S22 (Biochrom). OD_407nm_ was plotted against hemin concentration and data were fitted to a one-binding site model (with a calculated extinction coefficient of bound hemin of ε407=70 mM^−1^.cm^−1^) as described (9). Hemin binding affinity: tryptophane fluorescence quenching (emission from 300 nm to 400 nm; 280 nm excitation) was determined as a function of hemin concentration, using a Carry Eclipsefluorimeter. Curves Absorbance and fluorescence curves fitting were fitted to the one-binding site model with the following equation:

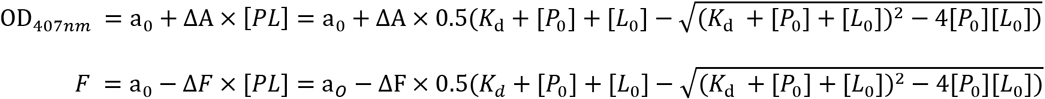

Here, OD_407nm_ and F are respectively the absorbance and fluorescence amplitudes. ΔA and ΔF are the normalized amplitude of the saturated absorbance and quenching, respectively, [PL] is the concentration of liganded protein, [P0] and [L0] are the concentrations of the total proteinand ligand, respectively and a0 is the starting absorbance/fluorescence. Fitting of these equation was was done with the non-linear regression function of GraphPad Prism 4 software.

#### EPR spectroscopy

X-band cw-EPR spectra were recorded with a Bruker Elexsys 500 X-band spectrometer equipped with a standard ER 4102 (Bruker) X-band resonator, a Bruker teslameter, an Oxford Instruments cryostat (ESR 900) and an Oxford ITC504 temperature controller. The spectra shown were recorded at 15 K with a modulation frequency equal to 100 kHz, a modulation amplitude equal to 25 gauss, a microwave power equal to 5 mW and a microwave frequency equal to 9.49 GHz.

#### *In vivo* imaging

Light emission from whole animals was measured in an *in vivo* imaging system (IVIS 200, Caliper Life Sciences, USA) equipped with the Living image software (version 4.0, Caliper Life Science, USA) as reported previously (4). Mice were anesthetized during imaging via inhalation of isoflurane. IVIS 200 was also used to evaluate luminescence on agar plates or from isolated organs (see above). Bioluminescence images were acquired with a 25 cm field of view (FOV), medium or large binning factor and an exposure time as indicated. A digital false-color photon emission image was generated according to photon counts within a constant region of interest (ROI) corresponding to the surface of the entire mouse. Rainbow images show the relative level of luminescence ranging from low (blue), to medium (green), to high (yellow/red). Photon emission was measured in radiance (p.s^−1^.cm^−2^ sr^−1^). Threshold parameters were chosen to maintain the luminescence detection under saturation level and were kept identical within an experiment. Images were adjusted for brightness and contrast using PhotoShop CS3 (Adobe Systems, San Jose, CA) with parameters kept identical in all images of the same figure.

## Suplemental Tables

**Table S1.**
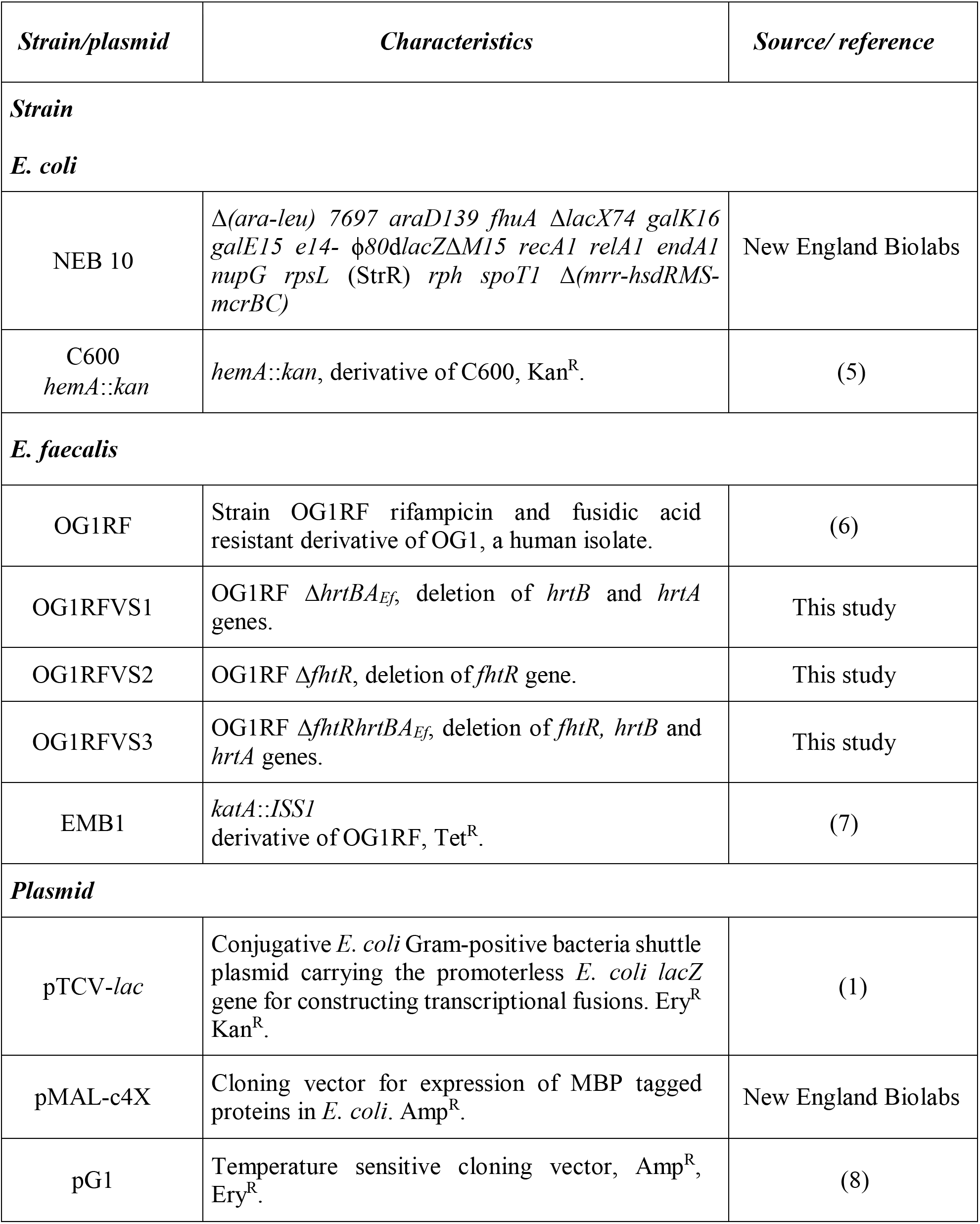
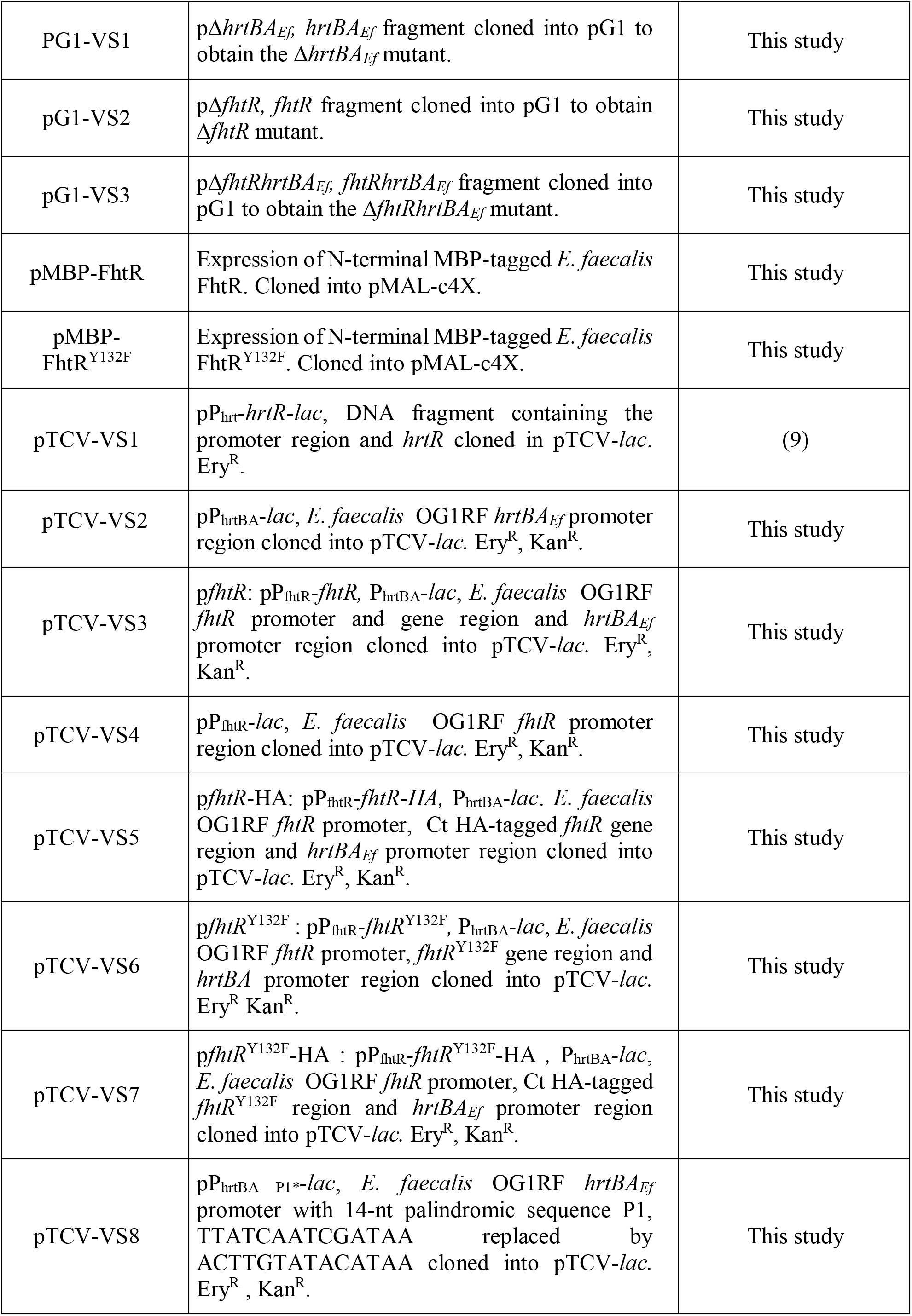
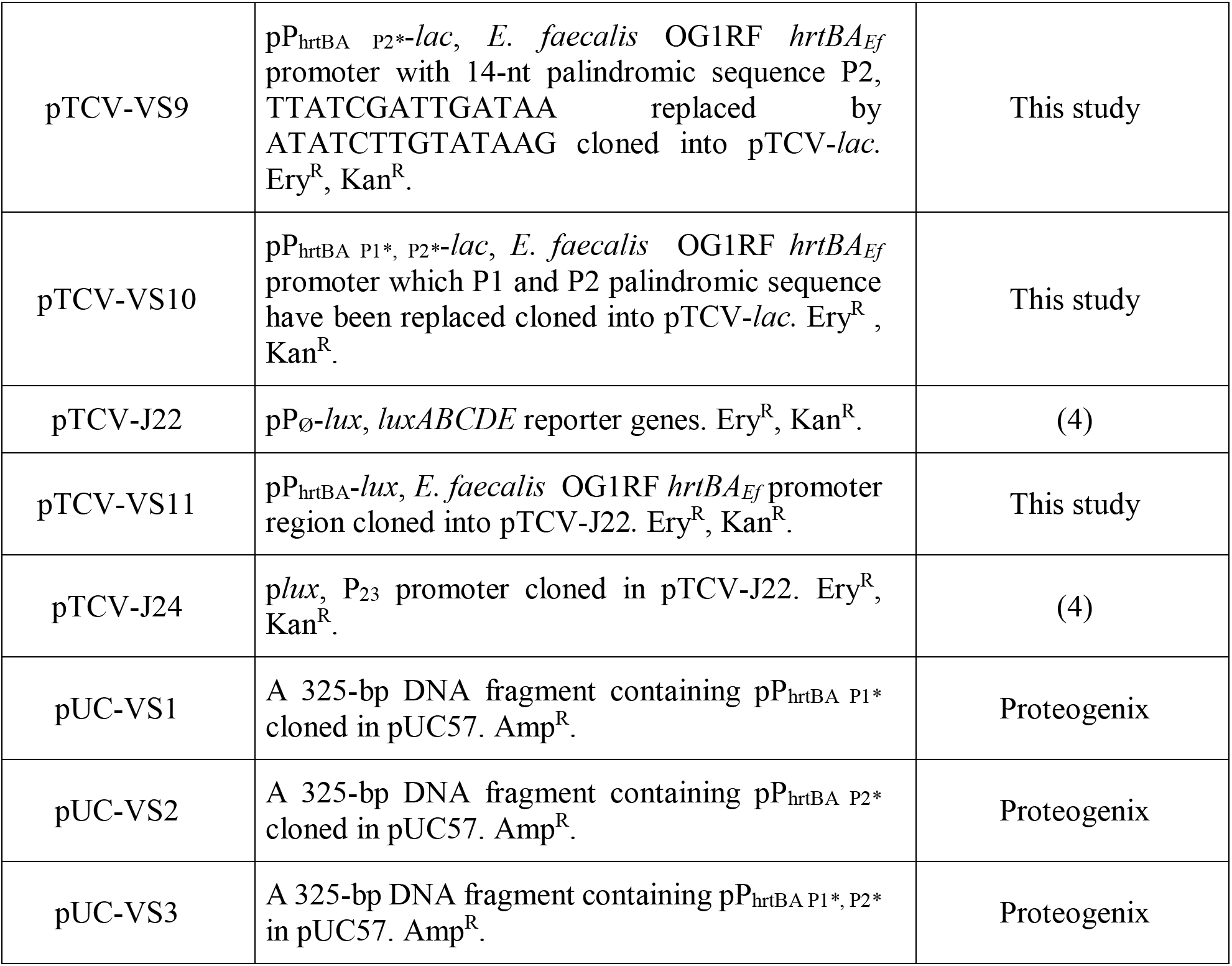
Strains and plasmids used in this study.

**Table S2.**
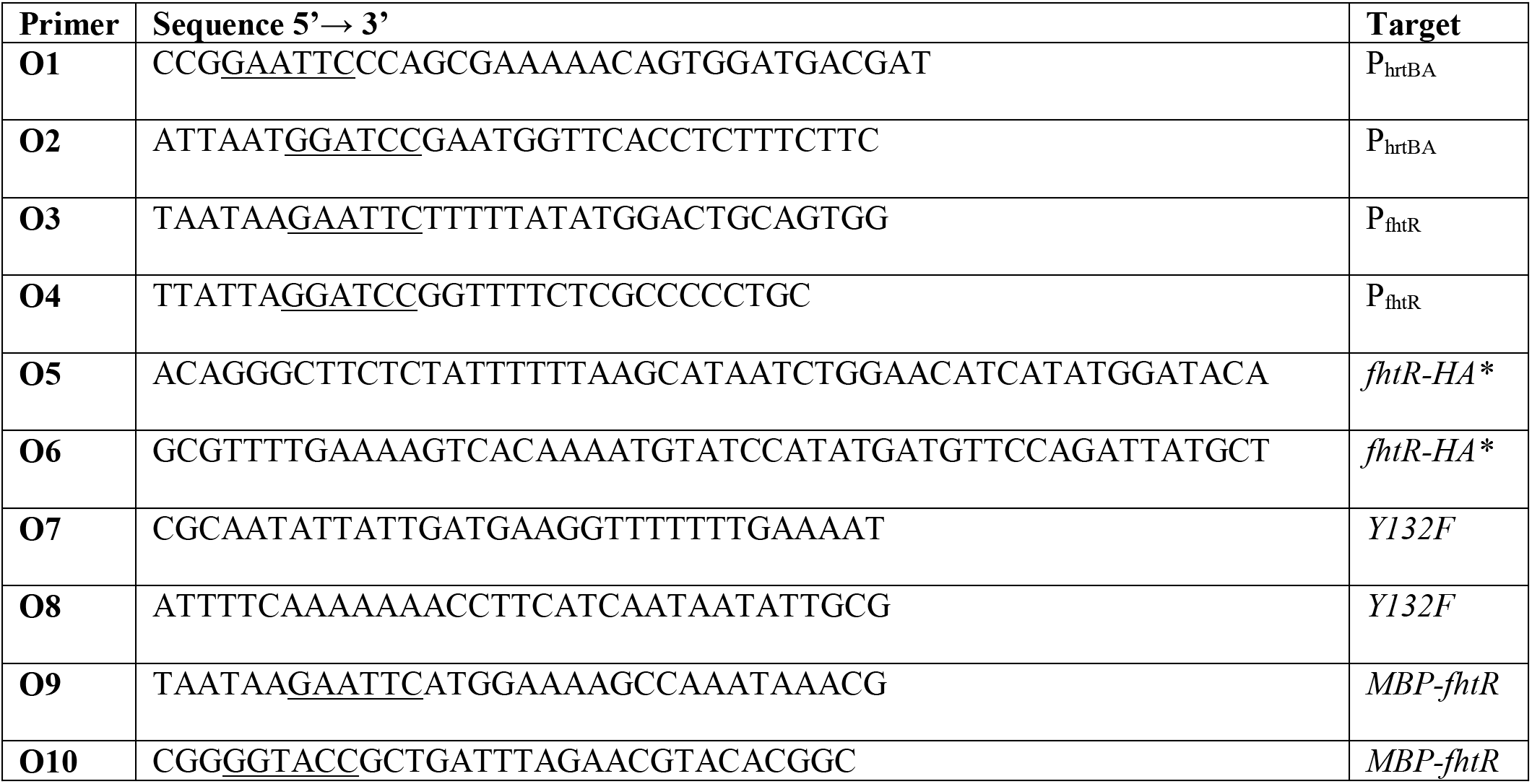
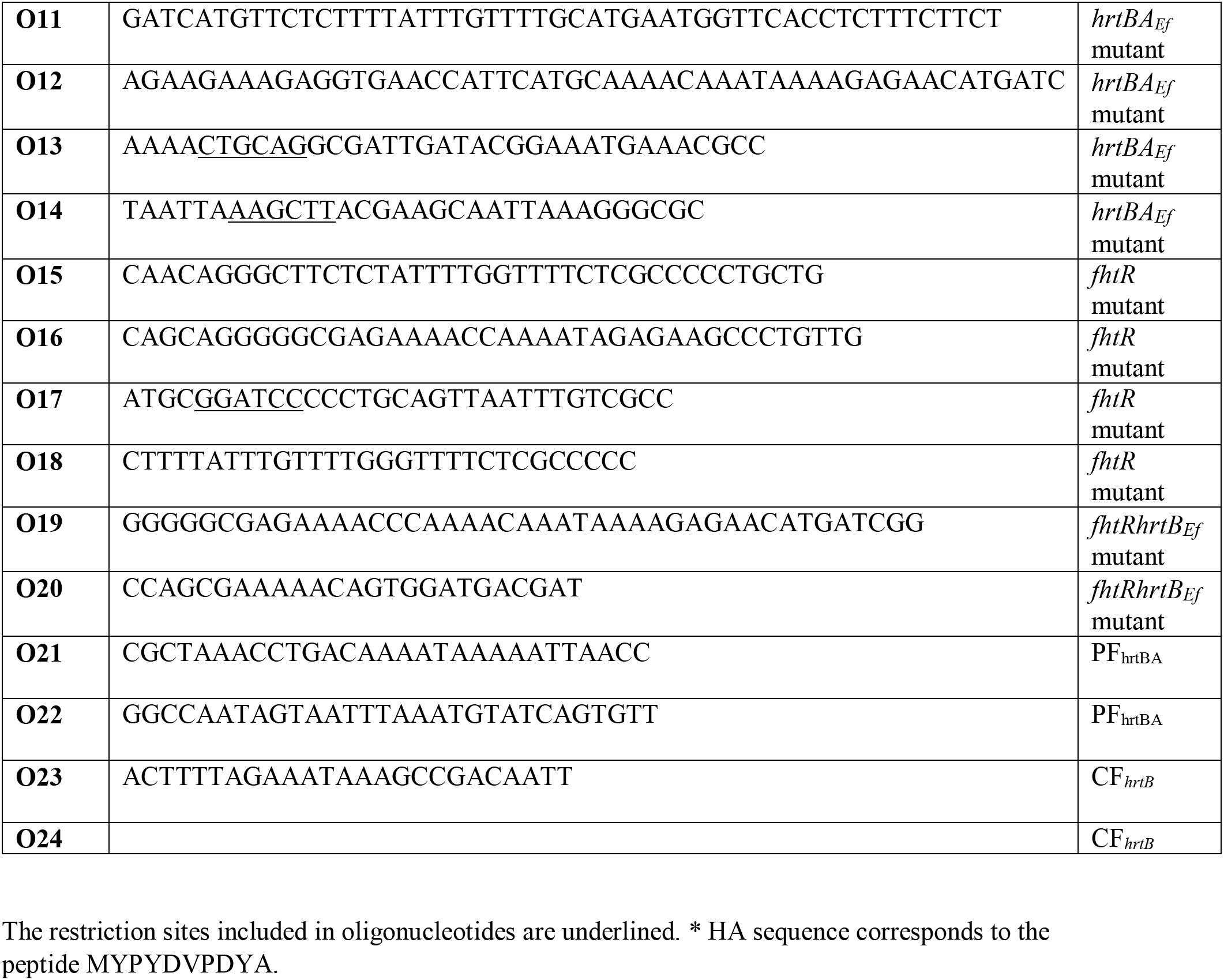
Oligonucleotides used in this study.

## Supplemental Figure Legends

**Figure S1.**
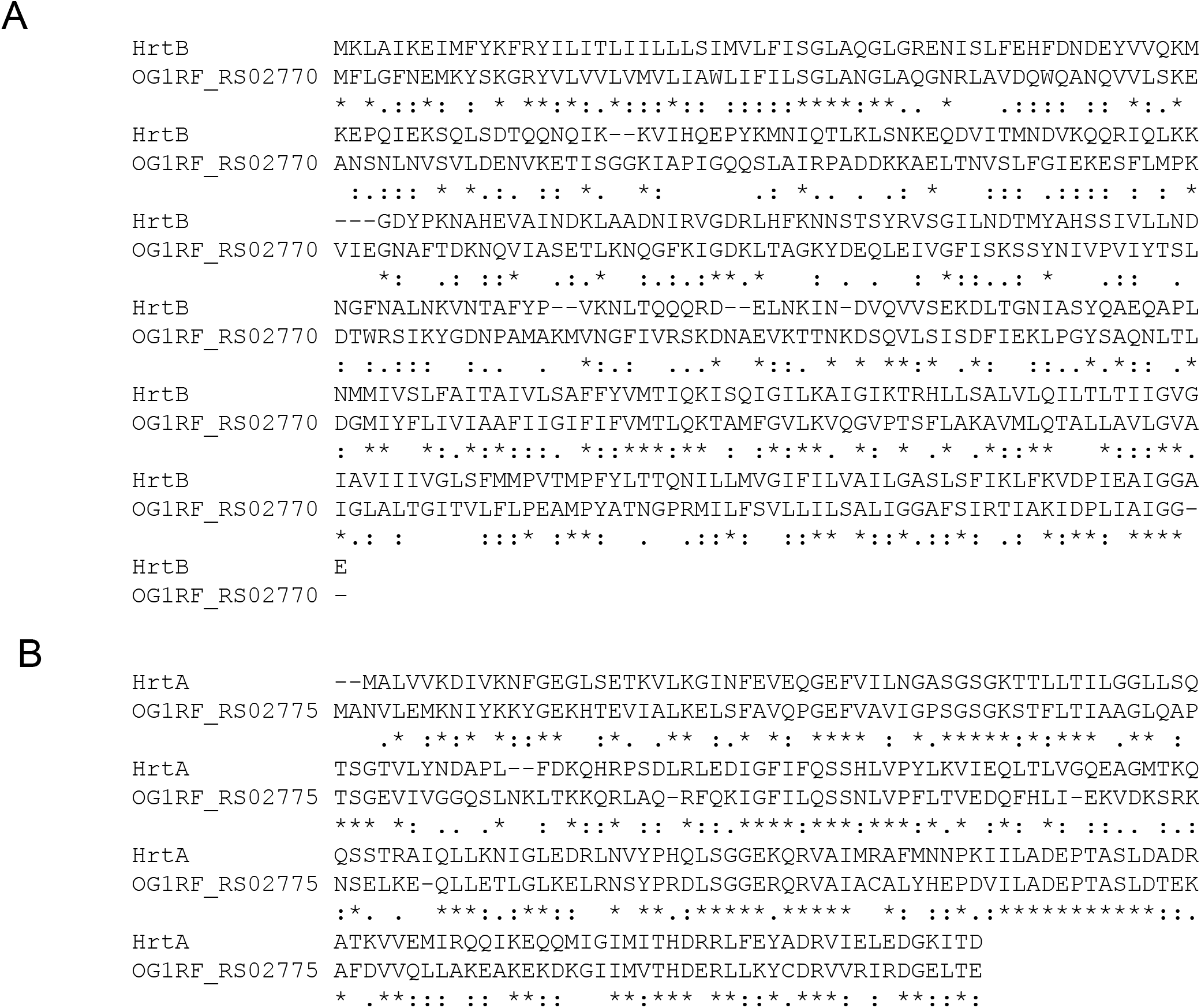
Alignement of OG1RF_RS02770 and OG1RF_RS02775 with HrtB_*Ef*_ (A) and HrtA_*Ef*_ (B) from *Staphylococcus aureus* respectively. AA sequences were aligned using Clustal W. Identical amino acids “*”; conserved amino acids “:”; partially conserved amino acids “.”.

**Figure S2.**
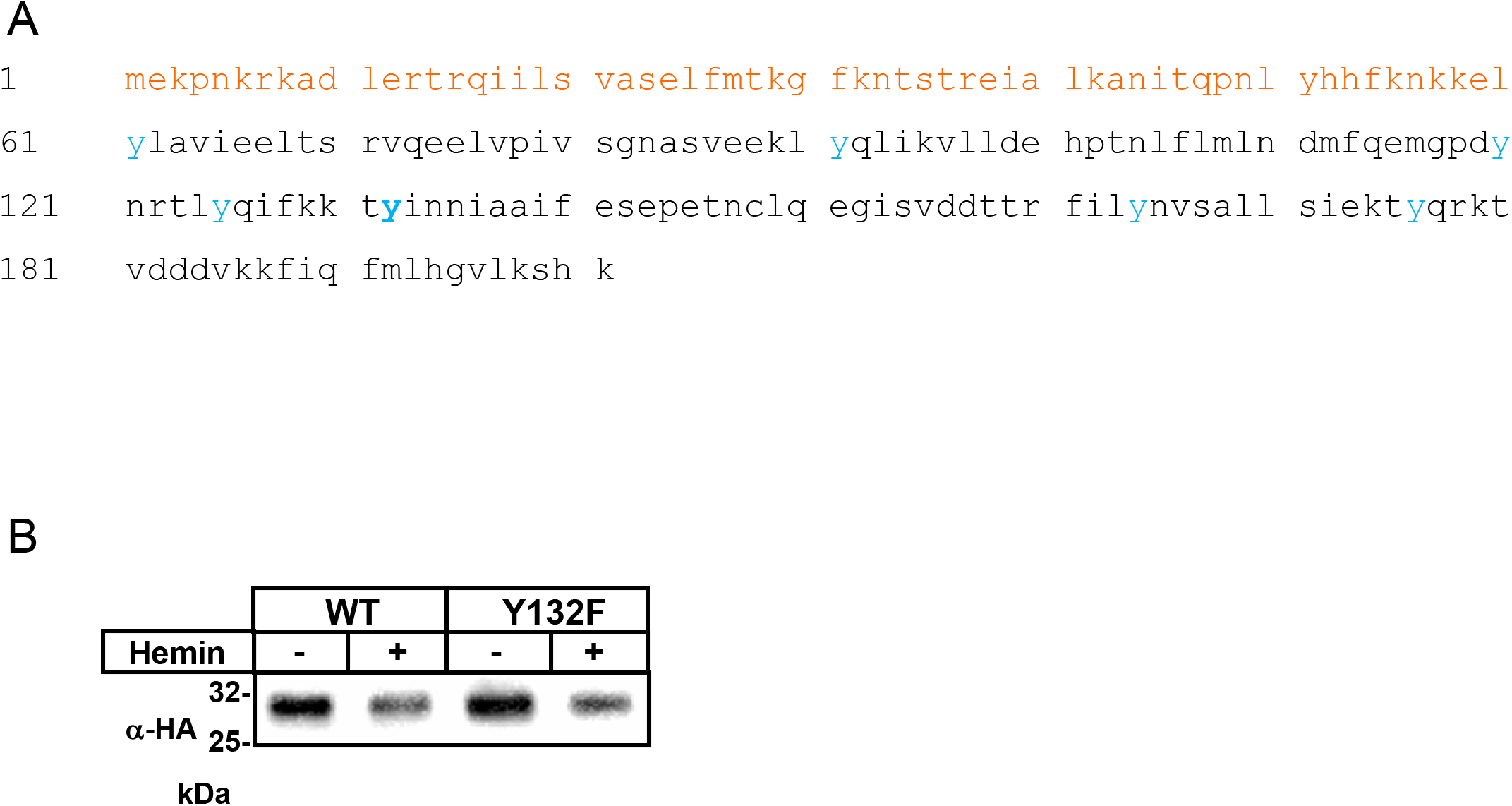
Hemin binding to FhtR implicates the tyrosine Y132. **(A) *E. faecalis* OG1RF FhtR AAs sequence.** Predicted Nt DNA binding region of the TetR family OG1RF_RS02765 ORF is shown in orange. Tyrosines are in blue and Y132 is in bold. **(B)** FhtR^Y132F^ is expressed similarly to WT FhtR. The Δ*fhtR* mutant was complemented either with p*fhtR*-HA or p*fhtR*^Y132F^-HA plasmids. FhtR and FhtR^Y132F^ expression were followed by Western Blot with a rabbit anti-HA antibody revealed by anti-rabbit antibody conjugated to peroxidase and detected by ECL. Bacteria were grown to OD_600nm_=0.5 and incubated with 2.5 μM hemin for 1.5 h. SDS-PAGE was performed on cell lysates (35 μg protein) per lane.

**Figure S3.**
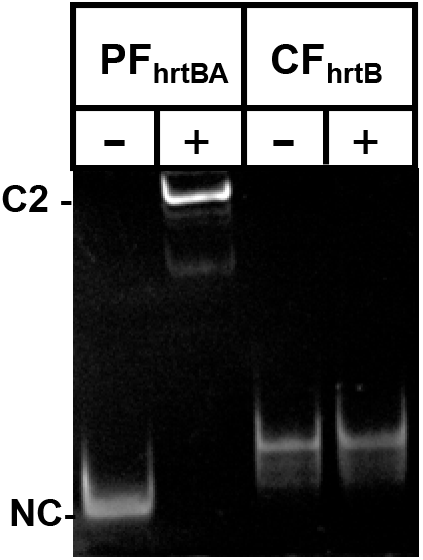
MBP-FhtR interaction with P_hrtBA_ is specific. 0.25 pmoles of the *hrtBA_Ef_* promoter fragment (FP_hrtBA_) or a similar size nt sequence in the coding region of *hrtB_Ef_* (CF_hrtB_) were incubated with 2.5 pmoles of MBP-FhtR. EMSA was performed as in Fig. 4 and DNA shift was visualized with GelRed (Biotium) following PAGE. C2 indicate the shifted DNA-protein complex and NC, the non-complexed DNA as in Fig. 4. Result is representative of 3 independent experiments.

**Figure S4.**
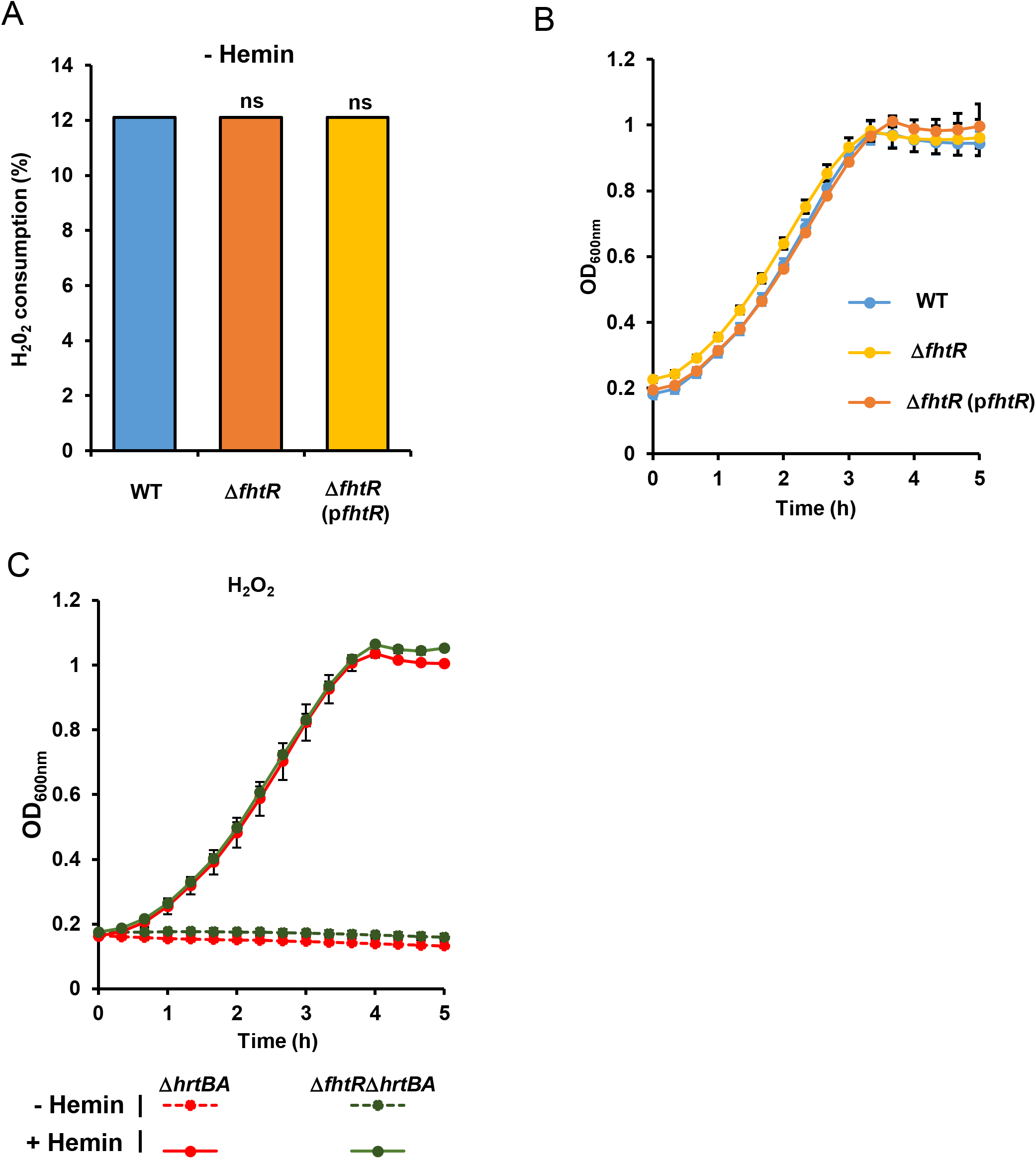
Control experiments for the study of KatA and FhtR interplay. **(A)** H_2_O_2_ background consumption in WT, Δ*fhtR* and Δ*fhtR*(p*fhtR*) grown in absence of hemin. Catalase activity was determined on equivalent number of bacteria from ON cultures incubated with 100 μM H_2_O_2_ for 30 min with the spectrophotometric FOX1 method based on ferrous oxidation in xylenol orange as in Fig. 6. Results expressed as the % of H_2_O_2_ concentration metabolized in respective strains and are the average ± S.D. of technical triplicates. ns, nonsignificant, oneway ANOVA followed by Dunnett’s posttest comparison to WT. Result is representative of 3 independent experiments. **(B)** Growth curves of WT, Δ*fhtR* and Δ*fhtR*(p*fhtR*). Overnight cultures were diluted to OD_600nm_ = 0.01 in M17G and grown to OD_600nm_ = 0.5. Cultures were distributed in a 96 well plate and incubated at 37 °C in a microplate Spark spectrophotometer (Tecan). OD_600nm_ was measured every 20 min. Results represent the average ± S.D. from triplicate technical samples. Result is representative of 3 independent experiments. **(C)** In absence of HrtBA_*Ef*_ expression, FhtR has no impact on KatA. Δ*hrtBA_Ef_* and Δ*fhtR*Δ*hrtBA_Ef_* strains were grown as in (B) with 1 μM hemin added at OD_600nm_ = 0.01. Results represent the average ± S.D. from triplicate technical samples. Result is representative of 3 independent experiments.

**Figure S5.**
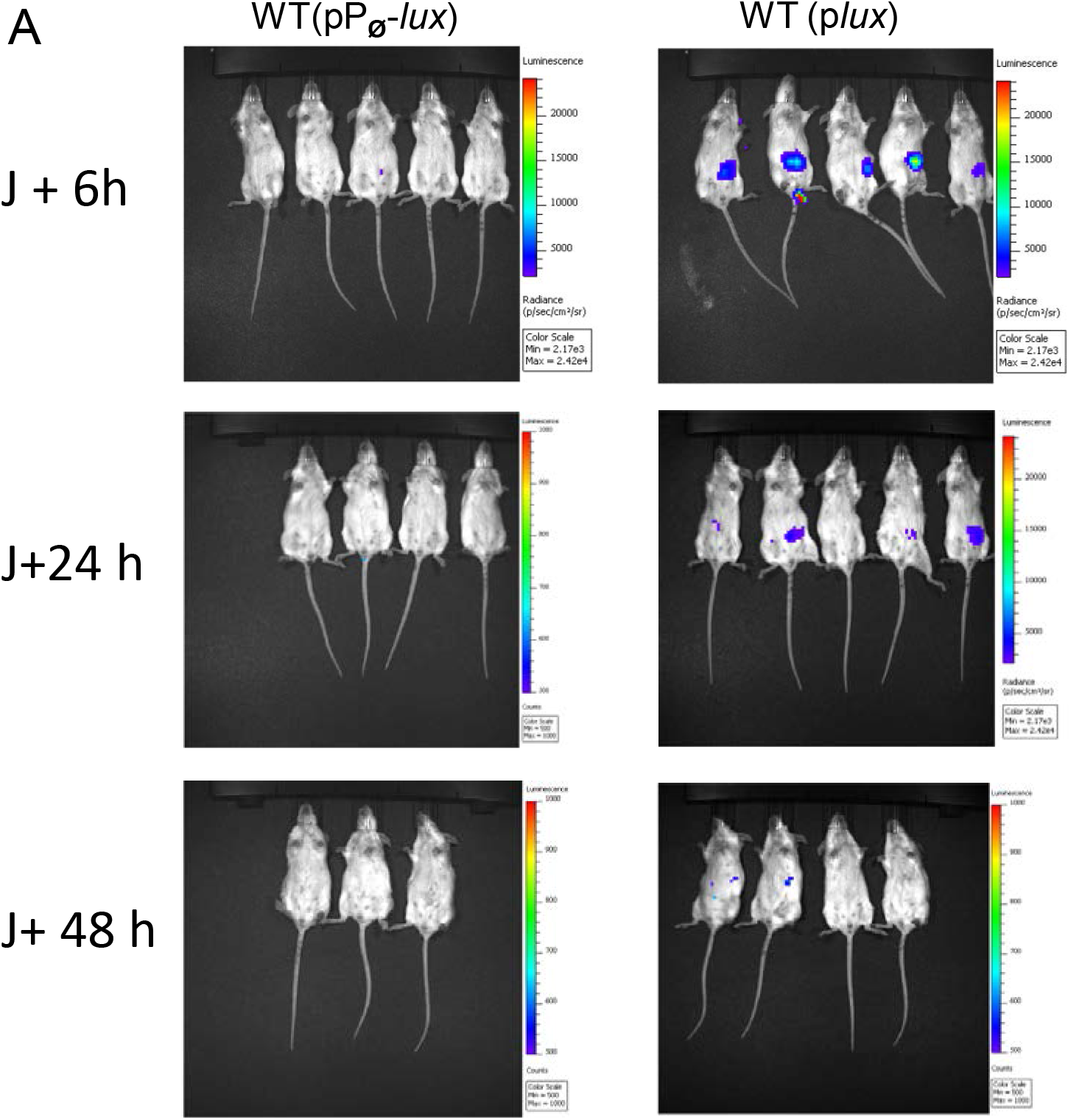
Time course of *Enterococcus faecalis OG1RF* survival in the GIT of mice following ingestion. Female BALB/c mice were force-feeded with 10^9^ CFU of WT(pP_∅_-*lux*) (control strain) or WT(p*lux*)(tracking strain). At the indicated times post-inoculation, mice were anesthetized with isoflurane and imaged in the IVIS 200 system (acquisition time, 20 min; binning 16).

**Figure S6.**
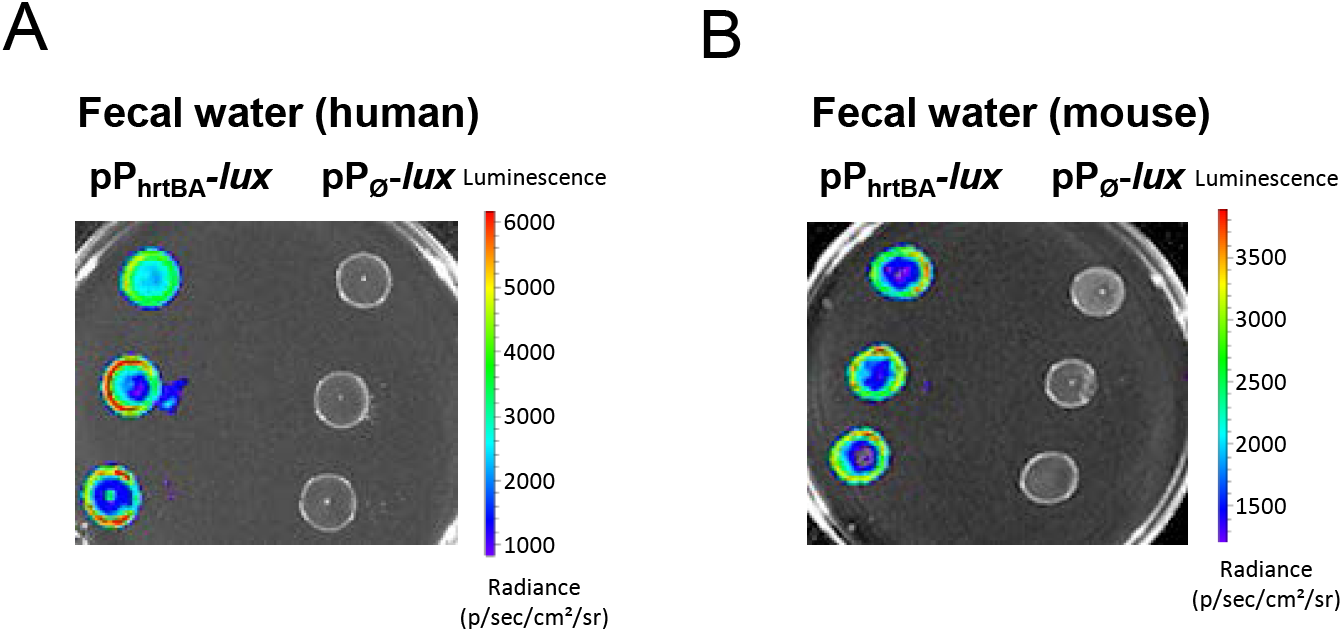
Human and mice fecal waters activate heme sensing. Human feces from 3 healthy human laboratory volunteers (A) and mice feces from 6 months old female BALB/c mice (B) were resuspended in PBS (25 μg/ml) and debris were removed by centrifugation at 5 000g at 4 °C for 30 min. Supernatants were sterile filtrated (0.22 μM) to remove bacteria from the extracts. Fecal water and WT OG1RF (pP_hrtBA_-*lux*) bacteria were plated as described in Fig. 8. Plates were imaged in the IVIS 200 system (acquisition time, 10 min; binning 8). Figure shows representative results corresponding to a total of 3 independent experiments.

**Figure S7.**
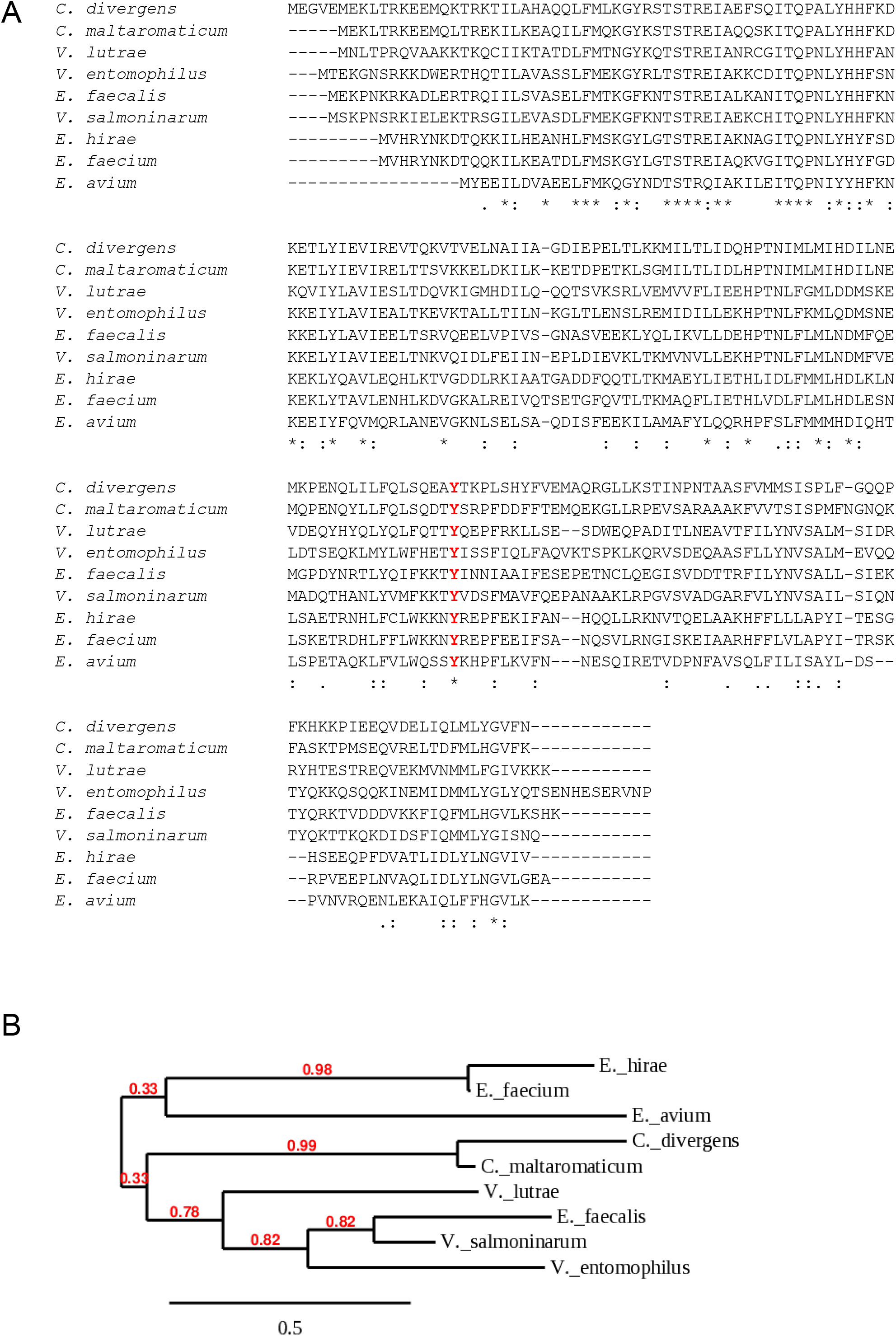
FhtR conservation across Gram positive bacteria. **(A)** FhtR orthologs across Gram positive bacteria. Blast analysis of FhtR shows that the protein is conserved in carnobacteri, enterococci and vagococci. FhtR AA sequences alignment from the following strains: *Carnobacterium divergens* (CDIV41), *Carnobacteriumc maltaromaticum* (DSM 20342 NODE_78), *Vagococcus lutrae* (CCUG39187), *Vagococcus entomophilus* (DSM 24756,1), *Enterococcus faecalis* (OG1RF), *Vagococcus salmoninarum* (NCFB 2777), *Enterococcus hirae* (ATCC9790), *Enterococcus faecium* (ATCC 8459), *Enterococcus avium* (ATCC14025)) were aligned using Clustal W. Identical amino acids “*”; conserved amino acids “:”; partially conserved amino acids “.”. **(B)** Phylogenetic analysis of FhtR amino-acid sequences. Predicted FhtR as in (A) were aligned using Clustal W, and evolutionary history was inferred using the neighbor-joining method. The tree is drawn to scale, with bootstrap values (× 1,000) shown above the nodes.

## Bibliography

1. Arias CA, Murray BE. 2012. The rise of the Enterococcus: beyond vancomycin resistance. Nat Rev Microbiol 10:266–78.

2. Agudelo Higuita NI, Huycke MM. 2014. Enterococcal Disease, Epidemiology, and Implications for Treatment. *In* Gilmore MS, Clewell DB, Ike Y, Shankar N (ed), Enterococci: From Commensals to Leading Causes of Drug Resistant Infection, Boston.

3. Mendes RE, Castanheira M, Farrell DJ, Flamm RK, Sader HS, Jones RN. 2016. Longitudinal (2001-14) analysis of enterococci and VRE causing invasive infections in European and US hospitals, including a contemporary (2010-13) analysis of oritavancin in vitro potency. J Antimicrob Chemother 71:3453–3458.

4. Van Tyne D, Gilmore MS. 2017. Raising the Alarmone: Within-Host Evolution of Antibiotic-Tolerant Enterococcus faecium. mBio 8.

5. Gruss A, Borezee-Durant E, Lechardeur D. 2012. Environmental heme utilization by heme-auxotrophic bacteria. Adv Microb Physiol 61:69–124.

6. Pishchany G, Skaar EP. 2012. Taste for blood: hemoglobin as a nutrient source for pathogens. PLoS Pathog 8:e1002535.

7. Kumar S, Bandyopadhyay U. 2005. Free heme toxicity and its detoxification systems in human. Toxicol Lett 157:175–88.

8. Anzaldi LL, Skaar EP. 2010. Overcoming the heme paradox: heme toxicity and tolerance in bacterial pathogens. Infect Immun 78:4977–89.

9. Frankenberg L, Brugna M, Hederstedt L. 2002. *Enterococcus faecalis* heme-dependent catalase. J Bacteriol 154:6351–6356.

10. Winstedt L, Frankenberg L, Hederstedt L, Von Wachenfeldt C. 2000. *Enterococcus faecalis* V583 contains a cytochrome bd-type respiratory oxidase. J Bacteriol 182:3863–3866.

11. Brugna M, Tasse L, Hederstedt L. 2010. In vivo production of catalase containing haem analogues. FEBS J 277:2663–72.

12. Hammer ND, Reniere ML, Cassat JE, Zhang Y, Hirsch AO, Indriati Hood M, Skaar EP. 2013. Two heme-dependent terminal oxidases power *Staphylococcus aureus* organ-specific colonization of the vertebrate host. MBio 4.

13. Joubert L, Dagieu JB, Fernandez A, Derre-Bobillot A, Borezee-Durant E, Fleurot I, Gruss A, Lechardeur D. 2017. Visualization of the role of host heme on the virulence of the heme auxotroph *Streptococcus agalactiae*. Sci Rep 7:40435.

14. Baureder M, Hederstedt L. 2013. Heme proteins in lactic acid bacteria. Adv Microb Physiol 62:1–43.

15. Fernandez A, Lechardeur D, Derre-Bobillot A, Couve E, Gaudu P, Gruss A. 2010. Two coregulated efflux transporters modulate intracellular heme and protoporphyrin IX availability in *Streptococcus agalactiae*. PLoS Pathog 6:e1000860.

16. Knippel RJ, Zackular JP, Moore JL, Celis AI, Weiss A, Washington MK, DuBois JL, Caprioli RM, Skaar EP. 2018. Heme sensing and detoxification by HatRT contributes to pathogenesis during *Clostridium difficile* infection. PLoS Pathog 14:e1007486.

17. Torres VJ, Stauff DL, Pishchany G, Bezbradica JS, Gordy LE, Iturregui J, Anderson KL, Dunman PM, Joyce S, Skaar EP. 2007. A *Staphylococcus aureus* regulatory system that responds to host heme and modulates virulence. Cell Host Microbe 1:109–19.

18. Lechardeur D, Cesselin B, Liebl U, Vos MH, Fernandez A, Brun C, Gruss A, Gaudu P. 2012. Discovery of an intracellular heme-binding protein, HrtR, that controls heme-efflux by the conserved HrtB HrtA transporter in *Lactococcus lactis*. J Biol Chem 287:4752–8.

19. Stauff DL, Skaar EP. 2009. *Bacillus anthracis* HssRS signalling to HrtAB regulates haem resistance during infection. Mol Microbiol 72:763–78.

20. Stauff DL, Skaar EP. 2009. The heme sensor system of *Staphylococcus aureus*. Contrib Microbiol 16:120–135.

21. Stauff DL, Torres VJ, Skaar EP. 2007. Signaling and DNA-binding activities of the *Staphylococcus aureus* HssR-HssS two-component system required for heme sensing. J Biol Chem 282:26111–21.

22. Joubert L, Derre-Bobillot A, Gaudu P, Gruss A, Lechardeur D. 2014. HrtBA and menaquinones control haem homeostasis in *Lactococcus lactis*. Mol Microbiol 93:823–33.

23. Ramos J, Martinez-Bueno M, Molina-Henares A, Teran W, Watanabe K, Zhang X, Gallegos M, Brennan R, Tobes R. 2005. The TetR family of transcriptional repressors. Microbiol Mol Biol Rev; 69:326–356.

24. Routh M, Su C-C, Zhang Q, Yu E. 2009. Structures of the AcrR and CmeR: Insight into the mechanisms of transcriptional repression and multi-drug recognition in the TetR family of regulatorsin the TetR family of regulators. Biochim Biophys Acta 1794:844–851.

25. Kumar R, Lovell S, Matsumura H, Battaile KP, Moenne-Loccoz P, Rivera M. 2013. The hemophore HasA from Yersinia pestis (HasAyp) coordinates hemin with a single residue, Tyr75, and with minimal conformational change. Biochemistry 52:2705–7.

26. Aki Y, Nagai M, Nagai Y, Imai K, Aki M, Sato A, Kubo M, Nagatomo S, Kitagawa T. 2010. Differences in coordination states of substituted tyrosine residues and quaternary structures among hemoglobin M probed by resonance Raman spectroscopy. J Biol Inorg Chem 15:147–58.

27. Halpern D, Gruss A. 2015. A sensitive bacterial-growth-based test reveals how intestinal Bacteroides meet their porphyrin requirement. BMC Microbiol 15:282.

28. Mimee M, Nadeau P, Hayward A, Carim S, Flanagan S, Jerger L, Collins J, McDonnell S, Swartwout R, Citorik RJ, Bulovic V, Langer R, Traverso G, Chandrakasan AP, Lu TK. 2018. An ingestible bacterial-electronic system to monitor gastrointestinal health. Science 360:915–918.

29. Fiorito V, Forni M, Silengo L, Altruda F, Tolosano E. 2015. Crucial Role of FLVCR1a in the Maintenance of Intestinal Heme Homeostasis. Antioxid Redox Signal 23:1410–23.

30. Rockey DC. 2010. Occult and obscure gastrointestinal bleeding: causes and clinical management. Nat Rev Gastroenterol Hepatol 7:265–79.

31. Seth EC, Taga ME. 2014. Nutrient cross-feeding in the microbial world. Front Microbiol 5:350.

32. Fiore E, Van Tyne D, Gilmore MS. 2019. Pathogenicity of Enterococci. Microbiol Spectr 7.

33. Brewitz HH, Hagelueken G, Imhof D. 2017. Structural and functional diversity of transient heme binding to bacterial proteins. Biochim Biophys Acta Gen Subj 1861:683–697.

34. Frankenberg N, Moser J, Jahn D. 2003. Bacterial heme biosynthesis and its biotechnological application. Appl Microbiol Biotechnol 63:115–27.

35. Maresso AW, Schneewind O. 2006. Iron acquisition and transport in *Staphylococcus aureus*. Biometals 19:193–203.

36. Price EE, Boyd JM. 2020. Genetic Regulation of Metal Ion Homeostasis in *Staphylococcus aureus*. Trends Microbiol doi:10.1016/j.tim.2020.04.004.

37. Orelle C, Mathieu K, Jault JM. 2019. Multidrug ABC transporters in bacteria. Res Microbiol 170:381–391.

38. Yang HB, Hou WT, Cheng MT, Jiang YL, Chen Y, Zhou CZ. 2018. Structure of a MacAB-like efflux pump from *Streptococcus pneumoniae*. Nat Commun 9:196.

39. Sharom FJ. 2006. Shedding light on drug transport: structure and function of the P-glycoprotein multidrug transporter (ABCB1). Biochem Cell Biol 84:979–92.

40. Bull-Henry K, Al-Kawas FH. 2013. Evaluation of occult gastrointestinal bleeding. Am Fam Physician 87:430–6.

41. Zeng MY, Inohara N, Nunez G. 2017. Mechanisms of inflammation-driven bacterial dysbiosis in the gut. Mucosal Immunol 10:18–26.

42. Berry EA, Trumpower BL. 1987. Simultaneous determination of hemes a, b, and c from pyridine hemochrome spectra. Anal Biochem 161:1–15.

43. Lechardeur D, Fernandez A, Robert B, Gaudu P, Trieu-Cuot P, Lamberet G, Gruss A. 2010. The 2-Cys peroxiredoxin alkyl hydroperoxide reductase c binds heme and participates in its intracellular availability in *Streptococcus agalactiae*. J Biol Chem 285:16032–41.

## Supplemental References

1. Poyart C, Trieu-Cuot P. 1997. A broad-host-range mobilizable shuttle vector for the construction of transcriptional fusions to B-galactosidase in Gram-positive bacteria. FEMS Microbiology Letters 156: 193–8.

2. Lechardeur D, Fernandez A, Robert B, Gaudu P, Trieu-Cuot P, Lamberet G, Gruss A. 2010. The 2-Cys peroxiredoxin alkyl hydroperoxide reductase c binds heme and participates in its intracellular availability in *Streptococcus agalactiae*. J Biol Chem 285:16032–41.

3. Maguin E, Prevost H, Ehrlich SD, Gruss A. 1996. Efficient insertional mutagenesis in lactococci and other gram-positive bacteria. J Bacteriol 178:931–5.

4. Joubert L, Dagieu JB, Fernandez A, Derre-Bobillot A, Borezee-Durant E, Fleurot I, Gruss A, Lechardeur D. 2017. Visualization of the role of host heme on the virulence of the heme auxotroph *Streptococcus agalactiae*. Sci Rep 7:40435.

5. Letoffe S, Delepelaire P, Wandersman C. 2006. The housekeeping dipeptide permease is the *Escherichia coli* heme transporter and functions with two optional peptide binding proteins. Proc Natl Acad Sci U S A 103:12891–6.

6. Dunny GM, Brown BL, Clewell DB. 1978. Induced cell aggregation and mating in *Streptococcus faecalis*: evidence for a bacterial sex pheromone. Proc Natl Acad Sci U S A 75:3479–83.

7. Baureder M, Hederstedt L. 2012. Genes important for catalase activity in *Enterococcus faecalis*. PLoS One 7:e36725.

8. Mistou MY, Dramsi S, Brega S, Poyart C, Trieu-Cuot P. 2009. Molecular dissection of the secA2 locus of group B Streptococcus reveals that glycosylation of the Srr1 LPXTG protein is required for full virulence. J Bacteriol 191:4195–206.

9. Lechardeur D, Cesselin B, Liebl U, Vos MH, Fernandez A, Brun C, Gruss A, Gaudu P. 2012. Discovery of an intracellular heme-binding protein, HrtR, that controls heme-efflux by the conserved HrtB HrtA transporter in *Lactococcus lactis*. J Biol Chem 287:4752–8.

